# Critical-like bistable dynamics in the resting-state human brain

**DOI:** 10.1101/2022.01.09.475554

**Authors:** Sheng H. Wang, Gabriele Arnulfo, Vladislav Myrov, Felix Siebenhühner, Lino Nobili, Michael Breakspear, Satu Palva, J. Matias Palva

## Abstract

Brain activity exhibits scale-free avalanche dynamics and power-law long-range temporal correlations (LRTCs) across the nervous system. This has been thought to reflect “brain criticality”, *i.e.*, brains operating near a critical phase transition between disorder and excessive order. Neuronal activity is, however, metabolically costly and may be constrained by activity-limiting mechanisms and resource depletion, which could make the phase transition discontinuous and bistable. Observations of bistability in awake human brain activity have nonetheless remained scarce and its functional significance unclear. First, using computational modelling where bistable synchronization dynamics emerged through local positive feedback, we found bistability to occur exclusively in a regime of critical-like dynamics. We then assessed bistability *in vivo* with resting-state magnetoencephalography and stereo-encephalography. Bistability was a robust characteristic of cortical oscillations throughout frequency bands from δ (3-7 Hz) to high-γ (100-225 Hz). As predicted by modelling, bistability and LRTCs were positively correlated. Importantly, while moderate levels of bistability were positively correlated with executive functioning, excessive bistability was associated with epileptic pathophysiology and predictive of local epileptogenicity. Critical bistability is thus a salient feature of spontaneous human brain dynamics in awake resting-state and is both functionally and clinically significant. These findings expand the framework of brain criticality and show that critical-like neuronal dynamics *in vivo* involves both continuous and discontinuous phase transitions in a frequency-, neuroanatomy-, and state-dependent manner.

## Introduction

Since Newton and Leibniz, differential equations have been used to describe natural phenomena that manifest *continuous* and smooth temporal evolution. Nonetheless, this classic approach fails in modelling many dynamics, particularly in biology and neuroscience, that show *discontinuity* and abrupt divergence into discrete states over time^1,2^. Catastrophic events emerging in complex systems, such as disasters in ecosystems or epileptic seizures in the brain, comprise an important subcategory of discontinuous phenomena and attract inter-disciplinary research to mitigate their detrimental consequences and to identify the underlying mechanisms^1–3^.

Neuronal population oscillations and their synchronization reflect rhythmical fluctuations in cortical excitability and regulate neuronal communication^4,5^. The “brain criticality hypothesis” posits that the brain, like many complex systems, operate near a “critical” point of a *continuous* transition^6^ between asynchronous and fully synchronous activity^7–10^. Operation near such a critical point endows the system with moderate mean synchronization, emergent power-law spatio-temporal correlations, and many functional benefits such as maximal dynamic range^11^, communication^12^, processing^13^, and representational capacity^14,15^. Conversely, inadequate or excessive synchrony are incompatible with healthy brain functions^6,16^ and represent coma-^17^ and seizure-like states^18^, respectively.

However, the classic brain criticality hypothesis does not offer an explanation to neuronal bistability, *i.e.,* discontinuous transitions between asynchronous and fully synchronous activity. Bistability *per se* is a well known phenomenon in neurophysiological dynamics and is salient, for example, in slow oscillations with up- and down-states observable across scales from intra-cellular^19,20^ to local-field potentials (LFP)^21,22^ in animal brains. In the human brain, while there are several lines of *in vivo* evidence for “critical-like” brain dynamics near a continuous phase transition^7,8,10,23^, evidence for discontinuous transitions, *i.e.,* bistable criticality, in awake resting-state brain dynamics has remained scarce.

Neuronal bistability in awake humans has only been reported by in a single series of electroencephalography (EEG) studies that reported bistable switching of alpha oscillations between putatively quiescent and a hyper-synchronized states^24–26^. Studies of whole-brain cortical activity in resting-state functional magnetic resonance imaging data (fMRI) also suggest spontaneous bistable switching between synchronous and asynchronous, or between integrated and segregated dynamics, respectively^27,28^. The underlying neuronal activity substrates at these multi-second time scales have, however, remained unclear.

Theoretical studies posit that a high degree of bistability is universally indicative of catastrophic shifts^1,2,29,30^. Hence, even if moderate bistability could characterize healthy brain dynamics, we hypothesize that high bistability in neuronal synchrony would be indicative of a shift from healthy to a pathological regime where neuronal populations abruptly jump between asynchronous and hyper-synchronized, seizure-prone states.

In this study, we asked whether awake resting-state human brain exhibits critical-like bistable dynamics. We first used generative modelling to establish how varying degree of bistable dynamics emerges as a consequence of introducing a slow positive local feedback^25^ that is conceptually equivalent to increasing demands for limited resources^31^. We then analyzed a large body of magnetoencephalography (MEG) and intracerebral stereo-EEG (SEEG) recordings of resting-state human brain activity. In both MEG and SEEG, we found that anatomically and spectrally widespread bistability characterized neuronal oscillations from δ (3-7 Hz) to high-γ (100-225 Hz) frequencies. In MEG, moderate resting-state bistability was correlated positively with executive functions. In SEEG, conversely, excessive resting-state bistability was co-localized with the epileptogenic zone and thereby associated with the pathophysiology underlying epilepsy. Bistable criticality thus constitutes a pervasive and functionally significant feature of awake resting-state brain dynamics.

## Results

### State-dependent noise induces bistable criticality *in silico*

To assess the emergence of bistability and its relationship with critical-like dynamics, we simulated a variant of the classic Kuramoto model, a simple generative model of synchronization dynamics^32^ (Supplementary Methods). Briefly, the conventional Kuramoto model has a single control parameter, ***κ***, that defines the coupling strength between neuronal oscillators. Higher ***κ*** leads to stronger synchrony among the oscillators that is typically quantified with “order”, *R*. Here, we introduced a second parameter ***ρ*** that scales state-dependent noise via local positive feedback in a manner that is conceptually equivalent to the state-dependency in the stochastic Hopf bifurcation^25,33,34^, or the slow resource-loading mechanisms leading to self-organized bistability^35^.

At small values of *ρ*, the model behaved similarly to a conventional critical-like system with a continuous second-order phase transition where a gradual increase of *κ* results in a monotonic increase of order (Fig 1A). At moderate order, *i.e.,* at the phase transition between low and high order, power-law long-range temporal correlations (LRTCs) ^36^ emerged in model order fluctuations and delineated a critical regime (Fig 1B). Here, LRTCs were quantified using the detrended fluctuation analysis (DFA) of the order time series (Supplementary Methods).

**Fig 1.**
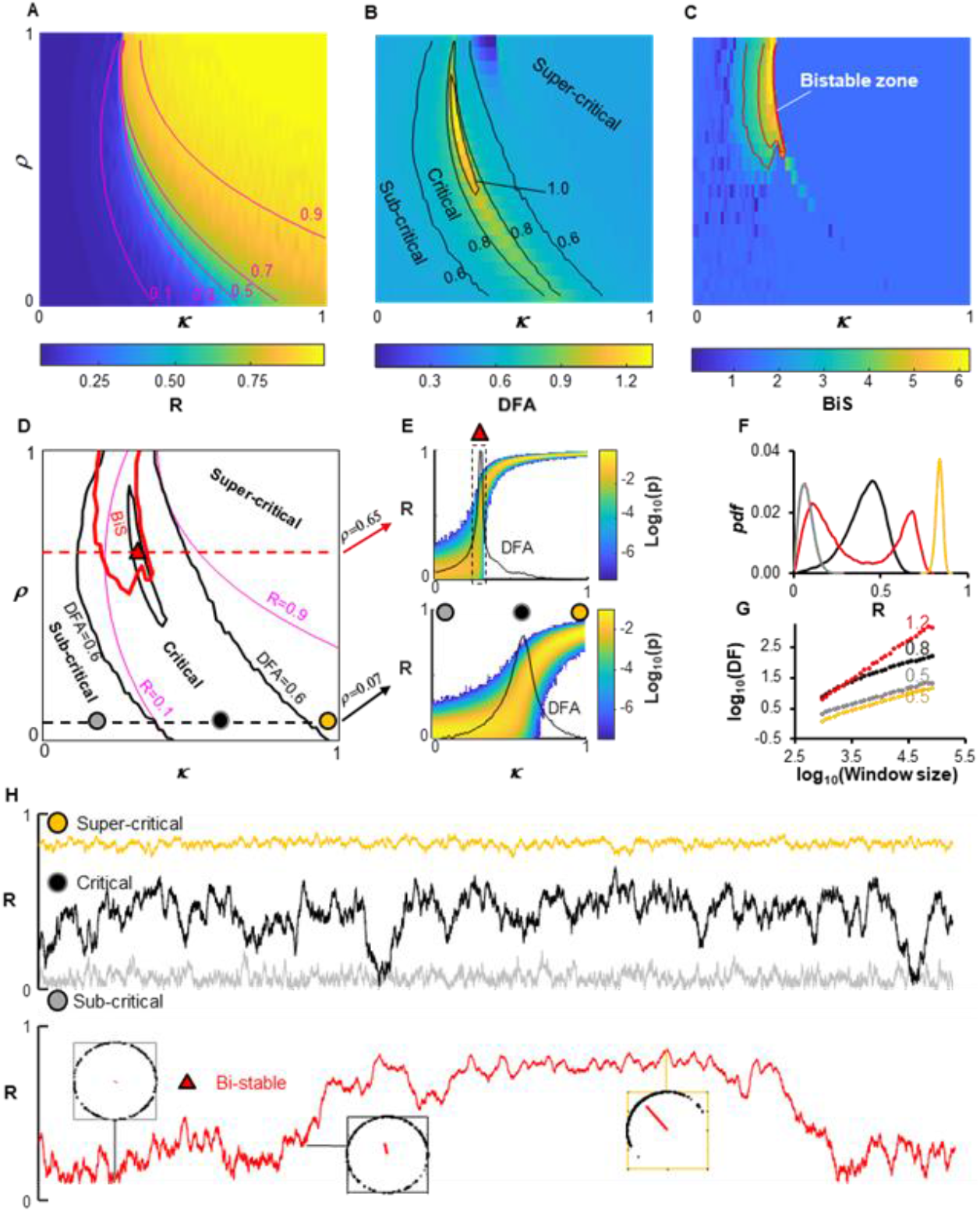
Bistability is caused by elevated state-dependent noise. (**A**) Kuramoto model order parameter (R), (**B**) Detrended fluctuation analysis exponent (DFA) – an estimate of LRTC, and (**C**) Bistability index (BiS) as functions of noise state-dependency (*ρ*) and the intrinsic control parameter (*a*). Each pixel is the mean of 50 independent model realizations. (**D**) Summary of overlapping regimes based on observation from (**B-D**), *i.e.,* the classic criticality is associated with small *κ* (black dashed line) whereas bistable criticality is caused by mid-to-high degree of *κ* (red dashed line); (**E**) Probability density of R in both normal and bistable criticality is in line with the Hopf bifurcation (see Supplementary). DFA peaks (black line) coincide with the phase transition. (**F**) Probability density (*pdf*) and (**G**) power-law scaling of the DFA fluctuation functions in classic and bistable critical regime marked in (**E**), colour coded. (**H**) Exemplary order parameter time series; insets in are the moments of Kuramoto oscillators (black dots) in low-, mid- and high-synch state (red vectors).

With increasing *ρ* values, the model synchronization dynamics became progressively bistable (Fig 1C) as evidenced by increasing values of the bistability index (BiS), an index of the relative fit of a bimodal versus a unimodal probability distribution *(pdf)* to the time series of squared order (*R^2^*, comparable to oscillation powers, see Supplementary). We found bistable dynamics exclusively within the critical regime (Fig. 1D). The presence of a bistable/discontinuous transition was also evident in the sudden increase in the order parameter at the critical value and the sharp peak in the DFA in contrast to the continuous transition (Fig 1E-F) and the representative time series (Fig 1H). Bistable dynamics at high *ρ* values thus likely reflect a first-rather than second-order phase transition. These *in silico* findings show that even in a minimal model, synchronization of oscillators may exhibit a continuum between classic and bistable critical dynamics under the influence of state-dependent noise via local feedback.

### Bistable criticality characterizes brain dynamics *in vivo*

We next assessed the presence of bistability and critical dynamics in meso- and macroscopic human brain activity in 10-minute resting-state recordings intracranially via stereo-electroencephalography (SEEG, *N* = 64) and source-reconstructed magnetoencephalography (MEG, *N* = 18), respectively. We first restricted analysis of the SEEG to neocortical grey matter contacts outside of the epileptogenic zone (EZ) (Fig 2-4). Although the anatomical sampling with SEEG is heterogenous across patients, the present cohort size yielded essentially a full coverage of the cerebral cortex (Supplementary Fig 2). We estimated LRTCs using DFA and bistability with BiS for narrow-band SEEG and MEG source amplitude time series that predominantly reflect local cortical synchronization dynamics.

**Fig 2.**
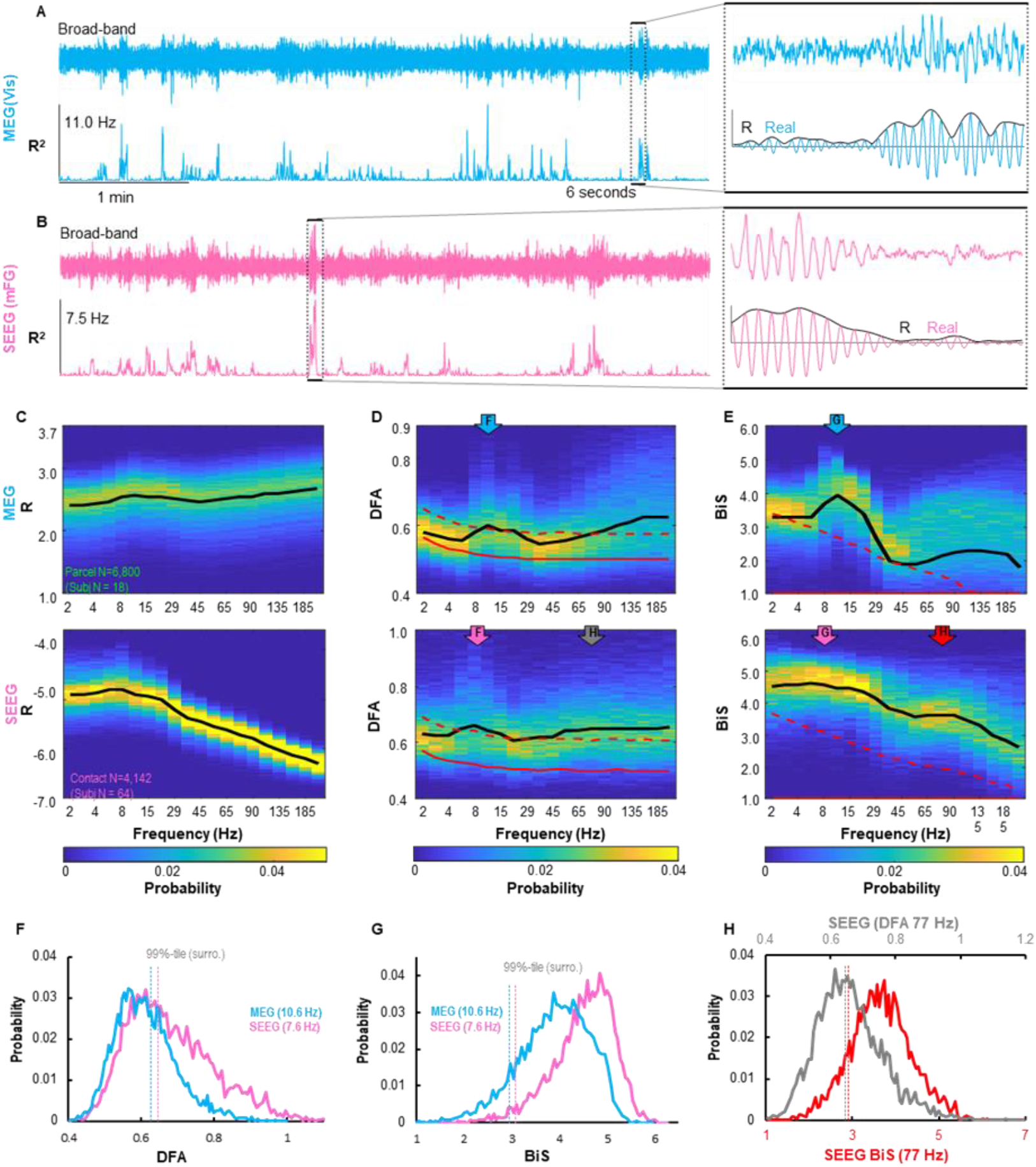
Bistability and LRTCs are robust, large-scale phenomena in MEG and SEEG. (**A**) Five minutes of broad band (top) and narrow-band filtered (11 Hz bottom) power (R^2^) time series from a MEG parcel located in visual area (Vis) in one subject; (**B**) Comparable output from five minutes of SEEG contact recorded from middle frontal gyrus (mFG) in one patient; insets: evidence of bistability as narrow-band traces switching between “up” and “down” states. (**C**) Group-level probability (z-axis) distribution of narrow-band (y-axis) mean amplitude (R), (**D**) DFA exponents, and (**E**) BiS estimates; data were pooled over all non-EZ SEEG contacts and MEG parcels; subject and contact/parcel number indicated in (**C**); black lines indicate mean of real data and red dashed lines are 99%-tile of surrogate observation. (**F-H**) Examples of narrow-band DFA and BiS probability distribution as indicated by colored arrows in (**D-E**).

**Fig 3.**
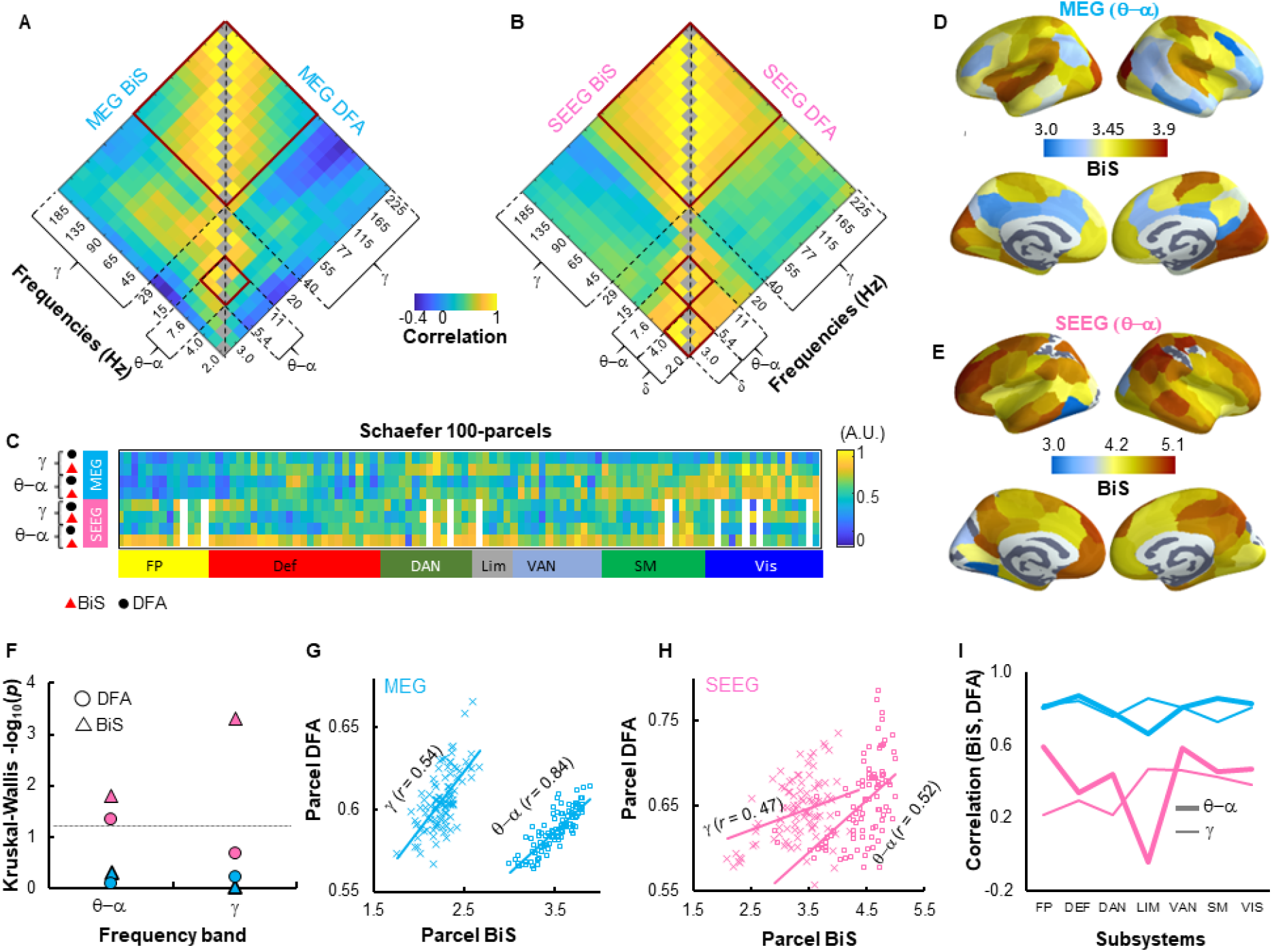
Bistability and LRTC were coexisting, correlated phenomena in MEG and SEEG. Neuroanatomical similarity (Spearman’s correlation) between group-average narrow-band BiS and DFA estimates of (**A**) MEG and (**B**) SEEG in Schaefer 100-parcel atlas; red boxes indicate frequency clusters showing high similarity. (**C**) Narrow-band group-averaged estimates were collapsed into θ–α (5.4-11Hz) and γ (40-225Hz) band based on similarity shown in (**A**); white-out columns in SEEG data indicate excluded parcels due to insufficient sampling (Supplementary). (**D**) Parcel-wise MEG group-average θ–α band BiS estimates presented in the cortex. (**E**) The same for SEEG group-average θ–α band BiS estimates. (**F**) Kruskal-Wallis one-way analysis of variance for group-level differences in DFA and BiS estimates between Yeo systems. Dashed line indicates –log_10_(*p value*) > 1.3, *i.e., p*<0.05. Correlations between group-average parcel BiS and DFA estimates in θ–α (cross) and γ band (circles) in (**G**) MEG and (**H**) SEEG, −log10(*p*) > 5, FDR corrected (Supplementary Fig 7). (**I**) Spearman’s correlations between within-subject-average BiS and DFA estimates in Yeo systems (subject N_MEG_ per system =18; N_SEEG_ per system = 50±9.4, range: 36-60, variable SEEG subject N per system due to heterogamous spatial sampling).

**Fig 4.**
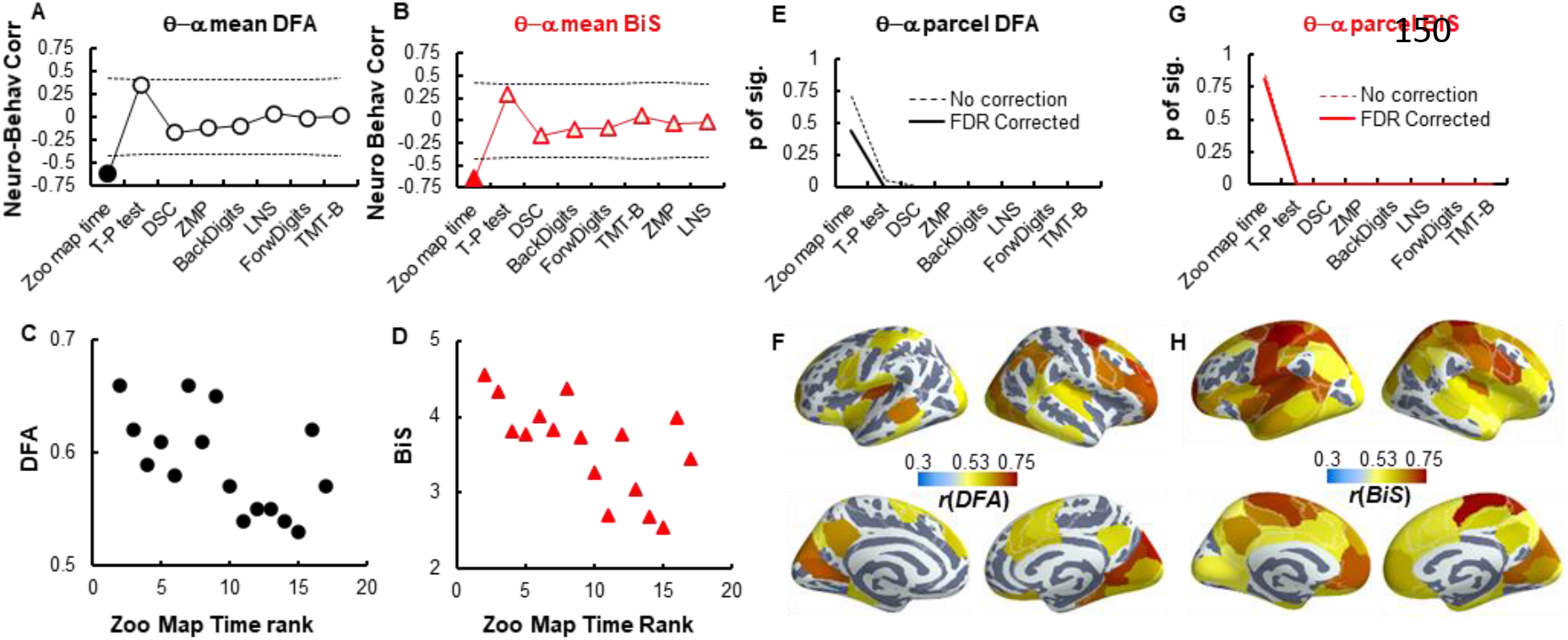
Executive functions were correlated with θ–α band DFA and BiS estimates in MEG subjects. (**A**) Spearman correlation between subject neuropsychological test scores and within subject mean parcel θ–α band DFA and (**B**) BiS estimates collapsed over parcels; dashed lines indicate 5% and 95%-tile of correlations for surrogate data (N_surrogate_ = 10^5^, FDR corrected, *p*-values see Supplementary Fig 10). (**C**) Scatter plots showing subject Zoo map time test and corresponding θ–α band parcel-collapsed DFA and (**D**) BiS estimates. Each marker in (**C-D**) stands for one subject. (**E**) Fraction of significant parcels that showed significant correlation between neuropsychological test scores and individual parcel θ–α band DFA and (**G**) BiS Estimates (*p*<0.05, FDR corrected, details see S.Fig 10). (**F**) Parcels showing significant correlations between Zoom map time scores and θ–α band DFA and (**H**) BiS estimates.

#### Bistability is anatomically widespread and spectrally prevalent

Visual inspection of narrow-band MEG and SEEG amplitude time series revealed salient examples of bistability as intermittent switching between low- and high-amplitude oscillations (Fig 2 A-B, for examples of model fitting for DFA and BiS estimates see Supplementary Fig 3). Statistical testing showed that both MEG-source signals and SEEG-electrode-contact LFP signals exhibited significant (*p* < 0.05, see Supplementary Fig 4) bistability and LRTCs across broad frequencies (Fig 2C-E). MEG showed a peak DFA and BiS estimates in the alpha (~11 Hz) frequency band whereas in SEEG, the BiS peak extended over δ (2-4 Hz), θ (4-7.6 Hz), and α (10-13 Hz) bands (Fig 2F-G). In SEEG, DFA and BiS estimates were overall stronger and occurred across more frequencies than in MEG (Fig 2E, F-H).

#### Neuroanatomical structure of bistability and LRTCs

We next characterized the neuroanatomical structure of bistability and inspected its anatomical relationship with LRTCs across frequencies. We first collapsed narrow-band BiS and DFA estimates of MEG parcels (400) and SEEG contacts into a standard atlas of 100 cortical parcels (see Supplementary Fig 4). Next, the neuroanatomical similarity within and between bistability and LRTCs were assessed by computing all-to-all Spearman’s correlations between narrow-band parcel BiS and DFA estimates.

Both MEG and SEEG showed high anatomical similarity between neighbouring frequencies. Correlations between slow and fast rhythms were negative in MEG (Fig 3 A) and weak in SEEG (Fig 3B). This indicates that regions tended to show bistability and criticality in a cluster of high or low frequencies, but not both. Based on these neuroanatomical similarities (red boxes, Fig 3 A-B, see also Supplementary Fig 5-6), we collapsed narrow-band BiS and DFA estimates into θ–α (5.4–11Hz) and γ-band (45-225 Hz) for further analyses (Fig 3C). The partitioning of β (15-30 Hz) band was not consistent and thus was not included (Supplementary Fig 5C).

MEG and SEEG cortical maps of θ–α band bistability revealed distinct neuroanatomical features. In MEG, visual (VIS), somatomotor (SM) and dorsal attention network (DAN) (Fig 3 C-D) exhibited greater BiS than expected by chance (*p*<0.05, two-tailed permutation test, 10^5^ permutations, not corrected for multiple comparisons, Supplementary Fig 9A-B). SEEG show high BiS in fronto-parietal (FP), ventral attention (VAN), default network (DEF), and limbic (LIM) systems (Fig 3C, E). Although comparable to the values in found in MEG, VIS showed the lowest BiS in SEEG (*p*<0.05, two-tailed permutation test, 10^5^ permutations, not corrected for multiple comparisons, Supplementary Fig 9C-D).

A Kruskal-Wallis test for variance among subjects’ BiS and DFA estimates within each Yeo system revealed that in SEEG, individuals showed different levels of BiS and DFA estimates between systems (Fig 3F) with bistability greater in DEF, FP, and LIM than in VIS and SM (unpaired t-test, p < 0.05, FDR corrected, Supplementary Fig9 C-D). There was no statistically significant regional variation in MEG data.

In both MEG and SEEG, group-average parcel bistability was correlated with LRTCs (Fig 3 G-H, see also Supplementary Fig 7). We validated this analysis in narrow-band frequencies and found the results to converge well (Supplementary Fig 6). To further validate this relationship, we averaged parcel BiS and DFA within subjects for each Yeo system and found that the subject BiS were indeed correlated with their DFA estimates on systems-level (Fig 3I, Supplementary Fig 8).

### Bistability is functionally significant in healthy subjects

We next asked whether bistability and LRTCs would predict individual differences in cognition. We assessed working memory, attention, and executive functions with neuropsychological tests (Methods). We averaged the BiS and DFA estimates across the cortical parcels to obtain four subject-specific neurophysiological estimates: DFA_θ–α_, DFA_γ_, BiS_θ–α_, and BiS_γ_ (Supplementary Fig 10B), and correlated these against neuropsychological test scores. We found that θ–α band BiS and DFA estimates were negatively correlated (*p* < 0.05, FDR corrected) with the Zoo Map Test Execution Time (Fig 4A-D, see also Supplementary Fig 10C-D). Thus the subjects with greater θ–α band bistability and stronger LRTCs executed faster in this flexible planning task, which is well in line with prior observations linking LRTCs with cognitive flexibility^37^.

To inspect the neuro-behavioural correlations in greater anatomical detail, we computed Spearman’s correlations between neuropsychological scores and individual parcel BiS and DFA estimates. A large fraction of the cortex showed significant neuro-behavioural correlations of θ–α band BiS and DFA estimates with the Zoo Map Time test but not with other neuropsychological scores (Fig 4 E and G, see also Supplementary Fig 10E-F). The correlations of Zoo Map Time test with DFA estimates were most pronounced in fronto-parietal, limbic, somatosensory and, and visual areas (Fig 4 F), whereas the correlations with BiS estimates were widespread across the cortex (Fig 4H).

### Excessive bistability characterizes the epileptogenic zone

Excessive bistability may predispose complex systems to catastrophic events^29,38^. Under the influence of strong state-dependent noise, our model demonstrated increased sensitivity to coupling strength (Fig 1E), which suggests that strong bistability could be an early sign of shift towards supercritical hypersynchronization events, *i.e.,* epileptic seizures. We thus asked whether bistability estimated from seizure-free, inter-ictal-activity-free resting-state SEEG recording could be informative about epileptic pathophysiology. In particular, we addressed whether bistability could delineate the epileptogenic zone (EZ) and dissociate EZ signals from signals in nEZ contacts that reflect more healthy forms of brain activity.

Representative time series (Fig 5A-B) showed that the EZ contacts did not show conspicuous epileptic inter-ictal events (IIE), and the sparse IIEs were removed from analysis where found (Supplementary Methods). Interestingly, elevated > 80 Hz bistability of the EZ contact was already a visually salient characteristic and stronger in EZ than in a nearby nEZ contact from the same region (DFA, bistability fitting see Supplementary Fig 3 F-J). We assessed bistability and LRTCs in narrow-bands frequencies at the group level for nEZ-versus EZ-electrode contacts (Fig 5C-D). Collapsing narrow-band DFA and BiS estimates into broader frequency bands revealed significant differences between nEZ- and EZ-electrode contacts in β- and γ-band BiS estimates with effect sizes of 0.5 and 0.65 (Cohen’s *d*), respectively (Fig 5E). There was also a difference between nEZ- and EZ-electrode contacts in the δ-band DFA exponent with a Cohen’s *d* of 0.2.

**Fig 5.**
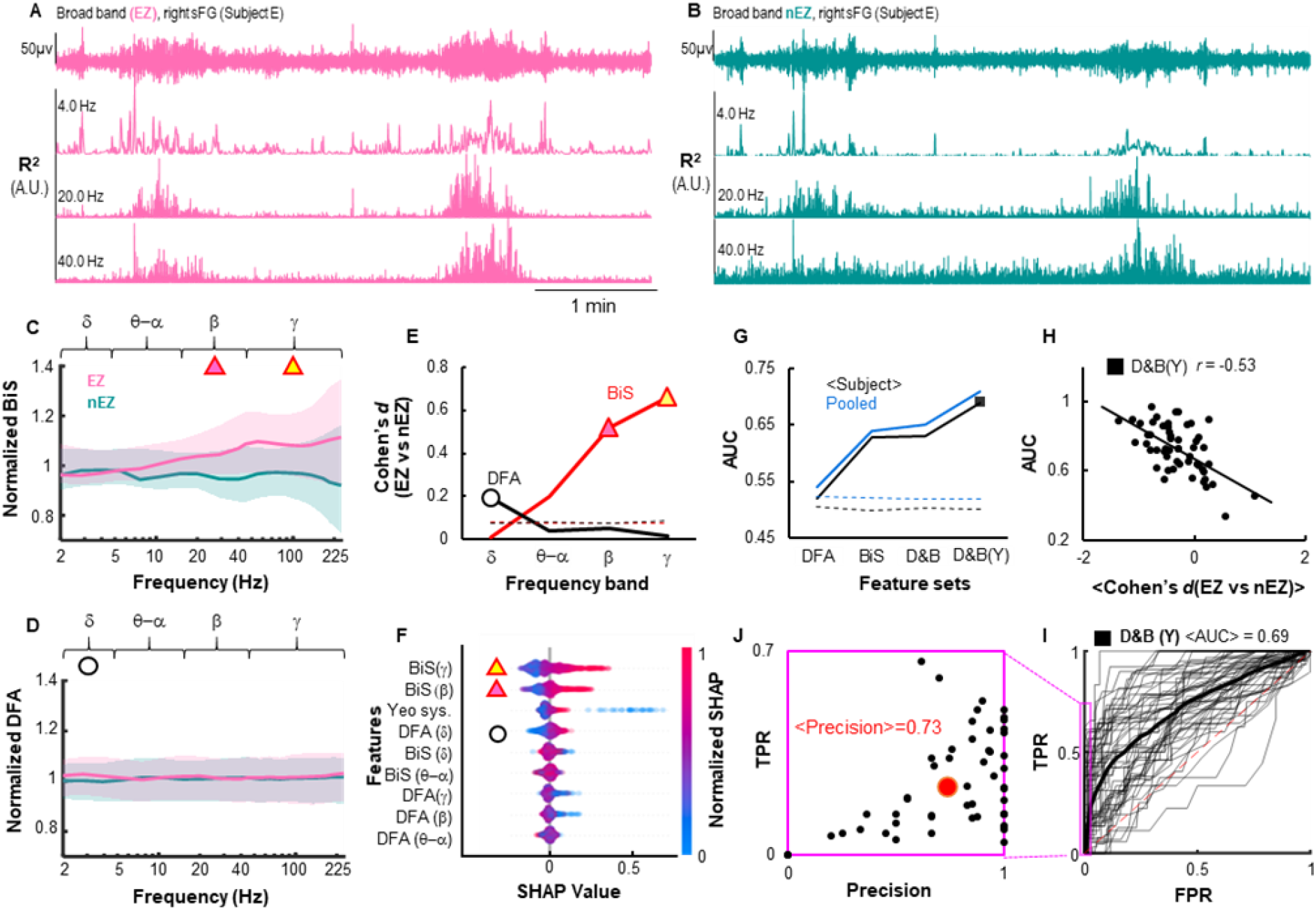
Bistability showed strong predictive power for pathophysiology. (**A-B**) Five minutes of broad-band traces and narrow-band power (R^2^) time series of an EZ (**A**) and a nEZ (**B**) cortical location recorded with two distinct electrode shafts in one subject. Both contacts were 19.7 mm apart within supervisor frontal gyrus (sFG) and were referenced with the same nearest white matter contact (Arnulfo et al., 2015). (**C**) Average normalized narrow-band BiS and (**D**) DFA estimates for all EZ (pink) and nEZ contacts (green). Shaded areas indicate 25% and 75%-tile. (**E**) The effect size of differences between EZ and nEZ contacts in frequency-collapsed BiS (red) and DFA (black). Dashed line indicates 99%-tile observation from surrogate data (N_surrogate_=1000). (**F**) Feature importance estimated using SHAP values. (**G**) The area under curve (AUC) of receiver operating characteristics (ROC) averaged across subjects (black) and the AUC of pooled within-subject classification results (blue) when using (*i*) DFA alone, (*ii*) BiS alone, (*iii*) combining DFA and BiS (D&B), and (*iv*) D&B plus contact loci in Yeo systems (D&B(Y)). Dashed lines indicate 99%-tile of AUC observed from 1,000 surrogates created independently for each of the four feature sets. (**H-J**) Post-hoc inspection of results derived using D&B(Y) feature set (the black marker in **G**). (**H**) Spearman’s correlation (*p* < 10^-6^, *n*=55) between individual AUC and within-subject mean Cohen’s *d* between EZ and nEZ in band-collapsed DFA and BiS. (**I**) Receiver operating characteristics of classification within subjects (thin lines) and mean ROC (thick). (**J**) Within-patient prediction precision as a function of TPR indicated by the magenta box from (**I**); the red marker indicates the population mean. *Precision* = *true positive* ÷ *reported positive.*

These group-level findings suggest that both bistability and LRTCs could constitute informative features for classifying nEZ- and EZ-electrode contacts. We thus conducted an EZ-vs-nEZ classification analysis using random forest algorithm^39^ and with frequency-collapsed BiS and DFA estimates as neuronal features, with the electrode contact location in Yeo systems as an additional feature. The cross-validation for the classification was a 80:20-partition (training:test) with 500 iterations. This revealed a reliable outcome with the area under curve (AUC) for the receiver operating characteristic reaching AUC = 0.8 ± 0.002 (mean ± std) (Supplementary Fig 11C). To identify the most informative components to the classifier, we assessed global and within-subject feature importance with the Shapley Additive exPlanations (SHAP) values^40^. The SHAP values corroborated that γ- and β-band BiS estimates were indeed the most important features, followed by contact location (Yeo system), and δ-band DFA (Fig 5F).

Given these encouraging group-level and contact-classification results, we quantified the within-subject accuracy of neuronal bistability in localizing the epileptogenic area. We used leave-one-out validation so that the EZ-vs-nEZ contact classification was performed for each patient with the rest of the patient serving as training data. Additionally, to independently evaluate the contributions of BiS and DFA estimates to classification accuracy, we implemented the classification with four feature sets: DFA alone, BiS alone, combining DFA and BiS (D&B), and combining D&B and SEEG contact location in Yeo systems (D&B(Y)).

Overall, the within-subject classification accuracy for EZ contacts was higher than chance level across all feature combinations (Fig 5G). Classification with all features yielded the best performance at an average AUC of 0.7±0.14 (Black marker, Fig 5G). BiS alone yielded a greater AUC than DFA alone. Including the contact-brain system as an additional feature to D&B increased the AUC by 0.06. The subject AUC values were correlated with the subject-specific differences in DFA and BiS estimates between EZ and nEZ (r=-0.53, p < 10^-6^) (Fig 5H) and were not affected by either the total numbers of contacts, EZ contacts, nor the ratio of EZ and nEZ contacts (Pearson’s correlation coefficient, *r* = −0.06, *p* = 0.66; *r* = −0.07, *p* = 0.61; *r* = −0.09, *p* = 0.50; respectively). Finally, the classifier yielded an average precision of 0.74 ± 0.30 (mean ± SD). While the true positive rate was 0.24 ± 0.17, the false-positive rate was only 0.03 ± 0.03 (Fig 5G and H), which shows that most EZ contacts identified with the bistability-based classification were correct even though the classifier did not identify all true EZ contacts.

## Discussion

We found here that bistable criticality is a pervasive characteristic of human brain activity and is both functionally significant in healthy brain dynamics and clinically informative as a putative pathophysiological mechanism in epilepsy. In a generative model of synchronization dynamics with positive feedback, bistability occurred exclusively within a critical-like regime. In both MEG and SEEG, we found significant bistability and LRTCs in spontaneous amplitude fluctuations of cortical oscillations widely across the neocortex and from δ- (2-4 Hz) to γ- (40-225 Hz) frequency bands. Moreover, as predicted by the model, bistability was positively correlated with LRTCs. As key evidence for functional significance, resting-state bistability was a trait-like predictor^41^ of healthy cognitive performance in MEG and clearly associated with epileptogenic pathophysiology in SEEG. These findings indicate that bistable criticality is an important novel facet of large-scale brain dynamics in both healthy and diseased human brain. Moreover, these observations strongly suggest that the brain criticality framework—currently focused on continuous phase transitions—should be expanded to include both continuous and discontinuous phase transitions (see Supplementary Theory).

In the model, we found bistability to occur exclusively within the critical regime so that the level of bistability increased monotonically with increasing positive feedback *ρ*. This state-dependent feedback thus acted as a continuous control parameter for shifting the model between a continuous and a discontinuous phase transition, at low or high levels of feedback, respectively. In both MEG and SEEG, band-collapsed θ–α (5.4-11 Hz) and γ (40-225 Hz) frequency cortical BiS and DFA estimates were correlated on group average level (see Fig 3 G-H) and among individuals within functional systems (see Fig 3I). These widespread correlations collectively suggest that a gradient from “low LRTCs and weakly bistable” to “high LRTCs and strongly bistable” is a systematic characteristic of the human brain dynamics.

A positive feedback loop is thought to be a generic mechanism^31,35,42–44^ leading to bistability in a wide range of modeled and real-world complex systems including the canonical sand-pile model^35^ and its variations^42^, ecosystems^29,45^, gene regulatory networks^25,46,47^, intra-cellular signaling^48,49^, and network models of spiking neurons^31,50^. In our model, the positive local feedback was implemented as state-dependent phase noise^25,34^. Three mechanisms have been proposed to account for feedback and state-dependency in microscopic neuronal dynamics^51^, whereas the exact neuronal mechanism for meso- and macroscopic state-dependency remains unclear. We postulate that the state-dependency be a slowly fluctuating physiological parameter conceivably reflecting the cortical excitability and corresponding resource demand^31,50,52^.

Bistable dynamics, in general, could be associated a dichotomy of both beneficial and detrimental outcomes. Organisms can operate in bistable mode that is thought to reflect a dynamic motif favourable to adaptation and survival^25,46,49,53^. However, a high degree of bistability characterizes many complex systems prone to catastrophic shifts^43,44^ such as sudden violent vibrations in aerodynamic systems^54^, irrevocable environmental changes^29,55,56^, wars and conflicts^57^. In healthy MEG subjects, BiS and DFA were correlated, and higher θ–α band BiS and DFA estimates predicted better cognitive performance. In the SEEG from epileptic patients, excessive β- and γ-band BiS, but not DFA, characterized EZ (see Fig 5E–F), which suggests a functional gradient wherein moderate bistability reflect functional advantages but high degree of bistability be a sign of pathological hyper-excitability; this pathological bistability could be associated with hyper-excitability, excessive synchrony, high resource demands, and likely subsequent oxidative stress and tissue damage^58^. This speculation is in accordance with biophysical models of seizures that suggest a crucial role of a discontinuous transition (a sub-critical Hopf bifurcation)^25,33^ in generalized seizures^52,53^.

With invasive SEEG, we found consistent and accurate performance of the BiS estimates in EZ localization, which suggests a great potential for broader clinical utility, *e.g.,* using non-invasive MEG or EEG. Future work could exploit the presence of widespread bistability to large-scale biophysical models of neural dynamics, building on the analytic link between the simplified model employed here and physiologically derived neural mass and mean field models. Whereas the simple model yields dynamical insights, the large-scale biophysical models are crucial for understanding biological mechanisms^33^, including those that describe seizure propagation in individual patient brain networks^59^.

## Materials and Methods

### The canonical Hopf bifurcation

The canonical model of sub- or supercritical bifurcation is ^25^:

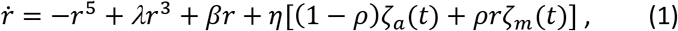

where 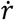 is the time derivative of a local neuronal activity *r* (a real number); *λ* is the shape parameter and *β* the bifurcation parameter; *η* scales the overall influence of noise; where *ζ_α_*(*t*) and *ζ_m_*(*t*) are additive and state-dependent noise respectively, and they are two uncorrelated Wiener processes; the parameter *ρ* weights the influence of state-dependent noise. Different combinations of *λ* and *β* result in either super-critical or sub-critical bifurcation (details in ^25^), which are associated with continuous or discontinuous (or second- or first-order) phase transition, respectively ^23,35,60^. When *r* described the amplitude of a two-dimensional system with phase 0, then (*eq.* 1) describes a Normal form stochastic Hopf bifurcation.

### The Kuramoto model

We studied first- and second-order phase transitions in a Kuramoto model with a modified noise term. The Kuramoto model is a generative model that can be used for studying the collective behaviours of a number of interconnected phase oscillators due to weak interactions ^32,61^. In a Kuramoto model, the dynamics of each oscillator *i* is a scalar phase time series *θ_i_* (0 ∈ 0:2*π*), coupled into a population ensemble ***θ*** as,

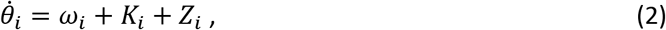

where, for any oscillator *i*, 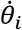 is the rotation of its phase *θ_i_*; *ω_i_* is the natural (uncoupled) frequency of *i*; *K_i_* = *K_i_*(***θ***) the coupling between *i* and the rest oscillators of the ensemble, and *Z_i_* is a stochastic term. The degree of synchrony of the ensemble (*i.e.*, order parameter or mean field) is the outcome of tripartite competition for controlling the collective behaviours of all oscillators: *ω_i_* and *Z_i_* are desynchronizing factors whereas *K_i_* is the synchronizing factor. Here, *ω_i_* follows a normal distribution with mean of zero (Hz), meaning without loss of generality, the system can be observed on a rotating phase plan with arbitrary angular velocity. In the classic model, the coupling term *K_i_* is defined as the *i*-th oscillator adjusts its phase according to interactions with all other oscillators in the system through a pair-wise phase interaction function:

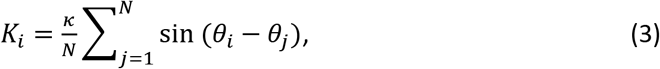

here, *κ* is a scalar number representing coupling strength, *N* =200 is the number of oscillators in the ensemble. For simplicity, here we used a fully coupled network to avoid other families of emerging dynamics due to nodal or network structural disorders, *e.g.,* Griffiths phase ^62,63^. In addition, we found that with a Gaussian nodal-weight distribution, the model behaved identically to the fully coupled networks. We modified the noise term in line with the Hopf bifurcation (*eq.* 1) as:

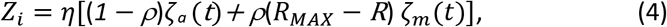

here, *ζ_a_*(*t*) and *ζ_m_*(*t*) are additive and multiplicative noise, respectively – as described in (*eq*. 1), however, in (*eq*. 4) *ζ_a_*(*t*) and *ζ_m_*(*t*) are uncorrelated and independent Gaussian phase noise with zero mean and unit variance; *ρ* scales the influence of *ζ_m_*(*t*); note that the 2 bracketed terms are offset (*i.e*., (1 – *ρ*) and *ρ*) such that their combined effect stays approximately constant in magnitude; *R_MAX_* is the maximal order the population can reach (*e.g.*, slightly below 1 due to the presence of noise) and *R* is the current mean field (a scalar) that quantifies the degree of synchrony of the population at time *t*,

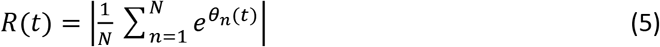

when viewing *R* from the complex phase plan, it essentially is the centroid vector of the population phase distribution: if the whole population is in full synchrony, *R*= *R_MAX_* →1; when there is no synchrony, *R* → 0 (see insets, Fig 1H).

### MEG recording and subjects

We recorded 10 minutes resting-state magnetoencephalographic (MEG) data from 18 subjects (11 males, 31.7±10.5, mean ± std, yeas of age) at the BioMag Laboratory, HUS Medical Imaging Center, Helsinki Finland. Subjects were seated in a dimly lit room and instructed to focus on a cross on the screen in front of them. Recordings were carried out at Meilahti hospital in Helsinki (detailed in Supp. Material). All subjects were screened for neurological conditions. The study protocol for MEG and MRI data obtained at the University of Helsinki was approved by the Coordinating Ethical Committee of Helsinki University Central Hospital (HUCH) (ID 290/13/03/2013) and was performed according to the Declaration of Helsinki.

We also assessed working memory, attention, and executive functions in these subjects with a battery of neuropsychological tests. These included (see Fig 4 x-axis): Zoo Map Time, Toulouse-Pieron test (TP), Digit Symbol Coding test, Zoo Map Plan, Digit Span forward and backward (BackDigits and ForwDigits, respectively), Letter-Number Sequencing (LNS), Trail Making Test parts A and B (TMT-A, TMT-B). Some subjects had missing/invalid behavioural scores, and we studied the neuro-behavioural correlations with dataset that at least had 16 valid subjects’ scores.

### SEEG recording and subjects

We acquired 10 minutes of uninterrupted, seizure-free resting-state brain activity with eyes closed from 64 drug resistant focal epilepsy patients (28 females, 30.1±9.1, mean±SD, yeas of age, see Supplementary Table 1) undergoing pre-surgical assessments (detailed in Supp. Material). The subjects gave written informed consent for participation in research studies. The study protocol for SEEG, computerized tomography (CT), and MRI data obtained in the La Niguarda Hospital were approved by the ethical committee of the Niguarda “Ca Granda” Hospital, Milan (ID 939), Italy, and was performed according to the Declaration of Helsinki.

Prior to surgery, medical doctors identified epileptogenic and seizure propagation zone by visual analysis of the SEEG traces ^64,65^. Epileptogenic areas (generators) are the region of interest that are necessary and sufficient for the origin and early organization of the epileptic activities ^66^. SEEG contacts recorded from such generators often show low voltage fast discharge or spike and wave events at seizure onset. Seizure propagation areas (receivers) are recruited during seizure propagation, but they do not initialize seizures ^59,67^. Contacts recorded from receivers show delayed, rhythmic modifications after seizure initiation in the generators. It is common to see regions demonstrating both generator and receiver dynamics, thus they were identified as generator-receiver. In this study, we refer to generator, receiver, and generator-receiver collectively as epileptogenic zone (EZ) to distinguish them from those that were tentatively identified as healthy non-EZ regions (nEZ).

## Abbreviations

BiS: bistability index
DFA: the scaling exponent obtained with detrended fluctuation analysis is an estimate of LTRCs
EZ: epileptogenic zone
nEZ: non-EZ, areas outside of the epileptogenic zone
*κ*: coupling strength between oscillators in the Kuramoto model
LRTCs: long-range temporal correlations
MEG: magnetoencephalography
*ρ*: the strength of the state-dependent noise in the Kuramoto model
R: the order parameter of the Kuramoto model
SEEG: stereo-EEG

## Acknowledgements

We would like to thank our Italian collaborators from the Centre of Epilepsy Surgery “C. Munari” at Niguarda Hospital, in particular Annalisa Rubino and Dr. Francesco Cardinale, for their support. We thank Drs. James Roberts for sharing code, and Serena di Santo, Stewart Heitmann, and Alexander Zhigalov for insightful discussions.

## Funding

The Academy of Finland SA 266402, 303933, and SA 325404 (SP)

The Academy of Finland SA 253130 and 296304 (JMP)

## Author contributions

Conceptualization: SHW, JMP

Funding acquisition: JMP, SP

Supervision: JMP

Methodology: SHW, MB, JMP

Software: SHW, MB, VM

Formal analysis: SHW

Resources: LN, GA

Data Curation: GA, FS

Visualization: SHW

Writing - Original Draft: SHW, JMP, MB, and SP

## Competing Interest Statement

All authors declare no competing interests. The funders had no role in study design, data acquisition, analyses, decision to publish, and preparation of the manuscript.

## Data and materials availability

Raw data and patient details cannot be shared due to Italian governing laws and Ethical Committee restrictions. Intermediate data, final processed data, and all code that support the findings of this study are available from the corresponding authors upon reasonable request.

## Supplementary Materials

### Supplementary methods

#### MEG data acquisition

A 306-channel MEG system (204 planar gradiometers and 102 magnetometers) with a Vectorview-Triux (Elekta-Neuromag) was used to record 10 minutes eyes-open resting-state brain activity from 18 healthy adult subjects at the BioMag Laboratory (HUS Medical Imaging Center, Helsinki Finland). For cortical surface reconstruction, T1-weighted anatomical MRI scans were obtained at a resolution of 1×1×1 mm with a 1.5 T MRI scanner (Siemens, Germany). This study was approved by the ethical committee of Helsinki University Central hospital and was performed according to the Declaration of Helsinki. Written informed consent was obtained from each subject prior to the experiment.

#### MEG data preprocessing and filtering

The Maxfilter software with temporal signal space separation (tSSS) (Elekta Neuromag Ltd., Finland) was used to suppress extra-cranial noise in sensors and to interpolate bad channels ^68^. Independent component analysis (Matlab Fieldtrip toolbox, http://fieldtrip.fcdonders.nl) was next used to identify and remove components that were correlated with ocular (identified using the EOG signal), heart-beat (identified using the magnetometer signal as a reference) or muscle artefacts ^69^. The FreeSurfer software (http://surfer.nmr.mgh.harvard.edu/) was used for subject MEG sources reconstruction, volumetric segmentation of MRI data, surface reconstruction, flattening, cortical parcellation, and neuroanatomical labeling with the Schaefer-2017 atlas ^70^. Each of the Schaefer-parcel belongs to a functional system ^71^ which informed later systems-level analysis.

The MNE software package was used to create head conductivity models and cortically constrained source models with 5000-7500 sources per subject and for the MEG-MRI co-registration and for the preparation of the forward and inverse operators ^72,73^. For each MEG subject, a cortical parcellation of 400 Schaefer-parcels was obtained using reconstruction-accuracy optimized source-vertex-to-parcel collapsing method ^74^. The broadband time series of these parcels were then filtered into narrow-band time series using a bank of 20 Morlet filters with *m* = 5 and log-linearly spaced center frequencies ranging from 2 to 225 Hz. For group-level analyses, subject DFA and BiS estimates were morphed from 400 Schaefer-parcels into 100 Schaefer-parcels.

#### SEEG data acquisition

Resting-state brain activity from 64 drug resistant focal epilepsy patients (28 females, 30.1±9.1, mean±SD, yeas of age, see S.Table 1) was acquired as monopolar local field potentials (LFPs) from brain tissue with platinum-iridium, multi-lead electrodes using a 192-channel SEEG amplifier system (NIHON-KOHDEN NEUROFAX-110) at 1 kHz sampling rate. Each penetrating electrode shaft has 8 to 15 contacts, and the contacts were 2 mm long, 0.8 mm thick and had an inter-contact border-to-border distance of 1.5 mm (DIXI medical, Besancon, France). The anatomical positions and amounts of electrodes varied according to surgical requirements ^64^. On average, each subject had 17 ± 3 (mean ± standard deviation) shafts (range 9-23) with a total of 153 ± 20 electrode contacts (range 122-184, left hemisphere: 66 ± 54, right hemisphere: 47 ± 55 contacts, grey-matter contacts: 110±25). The subjects gave written informed consent for participation in research studies and for publication of their data. This study was approved by the ethical committee (ID 939) of the Niguarda “Ca’ Granda” Hospital, Milan, and was performed according to the Declaration of Helsinki.

#### SEEG preprocessing and filtering

Cortical parcels were extracted from pre-surgically scanned T1 MRI 3D-FFE (used for surgical planning) using the Freesurfer package ^75^. A novel nearest-white-matter referencing scheme (its merits discussed in ^76^ was employed for referencing the monopolar SEEG LFP signals. An automated SEEG-electrode localization method was next used to assign each SEEG contact to a cortical parcel of Schaefer 100-parcel atlas with sub-millimeter accuracy ^77^. The SEEG electrodes were implanted to probe the suspected epileptogenic zones (EZ) while inevitably passing through healthy cortical structures. Contacts located at EZ are known to pick up frequent interictal spikes and generate abnormally large DFA ^78^. Therefore, EZ contacts and contacts recorded from subcortical regions such as thalamus, hippocampus and basal ganglia were excluded from analysis.

Nevertheless, inter-ictal events (IIE) such as spikes can be occasionally observed at non-EZ locations in some subjects during rests. These IIE are characterized by high-amplitude fast temporal dynamics as well as by widespread spatial diffusion, which need to be excluded to avoid bias to DFA and BiS estimates. We followed approach used in to identify such IIEs. Briefly, each SEEG contact broad-band signal was partitioned into non-overlapping windows of 500 ms in length; a window was tagged as “spiky” and discarded from LRTCs and bistability analyses when at least 3 consecutive samples exceeding 7 times the standard deviation above the channel mean amplitude. Last, narrow-band frequency amplitude time series was obtained by convoluting the broad-band SEEG contact time series with Morlet wavelets that were identical to that of MEG data.

#### Estimating LRTCs using detrend fluctuation analysis

LRTCs in 1D time series can be assessed with several metrics ^79^, and in this study, linear detrend fluctuation analysis (DFA) was used to assess specifically how fast the overall root mean square of local fluctuations grows with increasing sampling period ^36,80^. An estimated DFA exponent reflects the finite-size power-law scaling in narrow-band amplitude fluctuations based on the assumption that the gradual evolution of a mono-fractal process time series would result in a normal distribution where the fluctuation can be captured by the second order statistical moments such as variance ^81^. The computation of DFA briefly as follows (for rationales and technical details of the algorithm see ^82^):

1. The signal profile (***X***) of a signal was computed by computing the cumulative sum of a demeaned narrow-band amplitude of a MEG parcel or SEEG contact time series.
2. A vector of window widths (***T***) was defined in which the widths were linearly spaced on log10 scale between 10 and 90 seconds. The same scaling range was used across frequencies and for both MEG and SEEG, *i.e,* identical vector of ***T***. The lower boundary of 10 s was set to safely avoid high non-stationarity and the filter artifacts, *i.e.,* 20 cycles of the slowest rhythm of 2 Hz; the upper bound of 90 s was 15% of total sample of the resting-state recording.
3. For each window width *t*∈***T, X*** was partitioned into an array of temporal windows in which each window was of length *t* and with 25% overlap between windows *W*(*t*).

a. For each window *w*∈***W***(*t*), detrended signal ***W**_detrend_* was obtained by removing the linear trend, *i.e.*, subtracting the least-squares fit of samples of *w* from the samples of *w,* and then obtained the root mean square of ***W**_detrend_*(*RMS*(***W**_detrend_*)).
b. Finally, *F*(*t*), the detrended fluctuation of window size *t*, was obtained by computing the mean of RMS(***W**_detrend_*).
4. By repeating step (3) for all window lengths of ***T*** defined in step (2), ***F,*** a vector of *F*(*t*), *t* ∈ *T,* was obtained. The DFA exponent is the slope of the trend line of ***F*** as a function of ***T*** on log-log scale (Supplementary Fig 3 G, J).

#### Estimating bistability index (BiS)

The BiS index of a power time series (*R^2^*) derives from model comparison between a bimodal or unimodal fit of its probability distribution function (pdf); a large BiS means that the observed pdf is better described as bimodal, and when BiS → 0 the pdf is better described as unimodal. We followed the approach used in ^24,25^ to compute BiS. First, to find the pdf of power time series *R^2^,* the empirical *R^2^* was partitioned into 200 equal-distance bins and the number of observations in each bin was tallied. Next, maximum likelihood estimate (MLE) was used to fit a single-exponential function (*i.e.*, the square of a Gaussian process follows an exponential *pdf*):

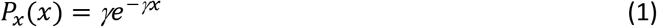

and a bi-exponential function:

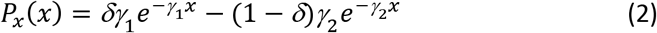

where *γ*_1_ and *γ*_2_ are two exponents and *δ* is the weighting factor.

Next, the Bayesian information criterion (BIC) was computed for the single- and bi-exponential fitting:

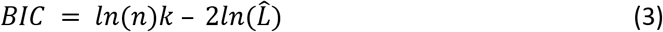

where *n* is the number of samples; 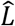 is the likelihood function; *k* is the number of parameters: *k* = 1 for single-exponential *BIC_Exp_* and *k* = 3 for bi-exponential model *BIC_biE_*. Thus BIC imposes a penalty to model complexity of the bi-exponential model ^83^ because it has two more degrees of freedom (second exponents and the weight δ) than the single exponential model.

Last, the BiS estimate is computed as the log10 transform of difference between the two BIC estimates as

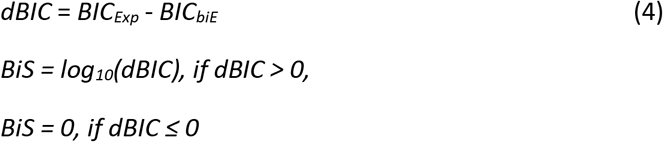

Thus, a better model yields a small BIC value, and BiS will be large if the bi-exponential model is a more likely model for the observed power time series (Supplementary Fig 3 F, I).

#### Constructing surrogate data

To determine the chance level observations of DFA and BiS in null hypothesis data, *i.e.,* without embedded non-linear critical-like structures but with the same power spectrum as the real physiological signals ^84^, phase randomized Fourier transform surrogates of the broad-band time series was constructed for each MEG parcel and SEEG contact, N_MEG_ = 6,800 and N_SEEG_ = 4,142 (Supplementary Fig. 4). The surrogate broad-band data were filtered into narrow-band data and their DFA and BiS estimates were subsequently computed. Thus, the real observation can be compared against the significance thresholds that were derived from the probability distribution of the surrogate data across frequencies (Fig. 2).

#### Morphing MEG and SEEG data into a standard atlas

The MEG and SEEG group-level analyses were conducted in a 100-parcel standardized Schaefer atlas ^70^. The group-level MEG data were obtained in two steps (top, Supplementary Fig 4). First, narrow-band DFA and BiS estimates were computed within subjects using a finer parcellation of 400-parcel, and the resulting estimates were morphed into 100-parcel within subjects by averaging children parcels; group level cortical maps were next obtained by collapsing subjects’ 100-parcel metrics.

Due to the variability in electrode location and other constraints in SEEG subjects, the morphing was done and verified differently. First, narrow-band DFA and BiS estimates of individual SEEG contacts were morphed directly into the Schafer 100-parcel atlas (bottom, Supplementary Fig 4). The resulting group-level parcel estimates were thus heterogeneous in terms of sampling. For example, one parcel may contain observations from a varying number of electrodes and/or subjects. Hence, the group-level estimate of each parcel was the median of all observations, and the estimate for each of the seven Yeo sub-systems was the median of its constituent parcels. Furthermore, only the parcels (n=90) sampled by at least 3 subjects and 10 SEEG contacts were kept for group-level analysis. The group mean parcel metrics (Supplementary Fig. 5B) were identical to that of raw data (overlay curves, Fig. 2D-E) thus confirming that heterogeneous SEEG sampling (Supplementary Fig. 2 and Supplementary Fig.4) did not bias the parcel-level metrics.

#### Clustering narrow-band frequency data

Frequency clustering analyses were conducted to reduce redundancy and for better interpretability of the narrow-band data (20 frequencies × DFA/BiS × 100 parcels). Spearman’s rank correlation coefficients were computed between the group-level DFA/BiS estimates of 100-parcel for all pairs of frequencies. This resulted in a set of 20 × 20 adjacency matrices *(**A**_f1,f2_*) representing the cortical topological similarity between frequencies. Next, the similarity matrix *A_f1,f2_* was partitioned using the unweighted pair group method with arithmetic mean algorithm ^85^, an agglomerative hierarchical clustering method, to obtain frequency clusters. The algorithm first builds a hierarchical tree through an iterative procedure to represent the distance between pairs of objects in *A_f1,f2_* (Supplementary Fig. 5C). In each iteration, two objects *p* and *q* with nearest distance *d*(*p, q*) were merged into a cluster, and *p* and *q* can be either an element from *A_f1,f2_* or a cluster of elements from *A_f1,f2_*; The distance function is defined as:

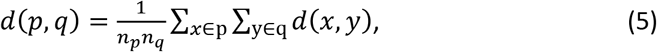

where *n_p_* and *n_q_* are the number of elements in *p* and *q* respectively, *d*(*x, y*) is the distance between *x*, *y*, and *x*, *y* are matrix elements from *A_f1,f2_*. The hierarchical tree was then used to partition the elements from *A_f1,f2_* into separated clusters, *e.g.*, if the height of *p* is close to the height of *q*, then their constituent elements are similar and therefore could be considered as a cluster (dashed boxes in Supplementary Fig. 5C, and solid boxes in Supplementary Fig. 5D).

#### Classifying pathophysiological SEEG contacts

The group-level frequency clustering analysis revealed that much of the narrow-band data were topologically correlated (Fig. 3). Hence, for the classification task, twenty narrow-band metrics were also collapsed into four frequency clusters as δ, θ–α, β, and γ band (Fig. 5A). As subjects varied greatly in their DFA and BiS estimates, band-collapsed data was normalized within subjects as [X-median(X))./max(X-median(X)], and thereby the differences between EZ and nEZ within subjects remained. The effect size of differences between band-collapsed and normalized DFA and BiS estimates were assessed with Cohen’s *d* and compared with the 99%-tile of Cohen’s *d* observed from 1,000 EZ-nEZ label-shuffled surrogate data (Fig. 5C).

The feature importance of these neuronal estimates were assessed with the SHapley Additive exPlanations (SHAP) values^40^. In addition to the neuronal scores, the contact location in Yeo systems was also included as an additional feature (Fig. 4D). The SHAP values is a generic metric to explain any tree-based model by explicating the local and global interpretability of features, which advances the transparency that conventional classifications approaches lack of. For solving the EZ-classification problem, the non-parametric random-forest method^86^ was employed. The random-forest algorithm is a machine learning method uses bootstrapped training dataset and combines the simplicity of decision trees with extended flexibility to handle new data^86^. The random-forest method allows multiple target class-labels (*e.g.*, nEZ plus three distinct EZ subtypes) over binary classifiers, and here the primary interest was to separate EZ and nEZ contacts.

### Supplementary materials

#### Theory: continuous and discontinuous phase transition in the brain

The classic Brain Criticality framework hypothesizes that, across the brain, neurons operate in a regime of continuous transitions between asynchronous and hyper-synchronous activity, which resembles the phase transition seen in numerous complex systems and is commonly known as the “critical point” or a “critical region”^7–10,23,87^. Recent theoretical research suggests that the brains could rather be a “quasi-critical” than a “true critical” system^50,88–90^. True criticality arises in idealized systems where energy is conserved and, with a small and constant drive, the system self-organizes into dynamics with one critical point^91^. True criticality is characterized by a continuous (second-order) phase transition between disorder and order^35^. At the phenomenological level, the ensuing critical dynamics are characterized by stationary fractional-Gaussian statistics^79^ and the emergence of spatial ^92^, temporal ^93^, and spatio-temporal power-law behaviour^88,91^.

On the other hand, quasi-critical systems^42,87,94^ differ categorically from the true critical systems by energy dissipation, as exemplified by forest fires and earthquakes^95,96^. The loss of energy in quasi-critical systems requires a “loading” mechanism to keep them from becoming quiescent. Simulations suggest that quasi-critical systems require external fine tuning to operate near the critical point^89,97^. The brain is energetically expensive, accounting for ~20% of the human energy consumption in adults^98^ and of up to ~66% in children^99^ with many physiological mechanisms serving energy replenishment and metabolic regulation^100^. In addition to glucose metabolism *per se,* the notion of “loading” mechanisms also encompasses resources and mechanisms that limit neuronal activity levels, for example synaptic vesicle depletion^101^ and post-synaptic depression^102,103^, respectively.

Theoretical studies show that neuronal systems with resource-consuming activity and slow resource loading^19,31,50,90^ may indeed exhibit dynamics with a discontinuous (first-order) phase transition when resource demands exceed the loading capacity. This gives rise to spontaneous neuronal *bistability*^50,90^. For example, in computational models of local populations operating near a marginally stable critical point^34^, excitatory neurons exhibit bistable firing rates when resource demands are high^50,90^. Likewise, in networks of such populations, the balance of resource depletion and recovery^102,103^ determines the switching between continuous and bistable transitions^31^.

#### Slow state-dependent noise ρ controls fast mean field

When investigating the behaviours in Kuramoto model (Fig 1), *ρ* and *κ* were held as constant values (*eq.* 4), which reflect that both variables fluctuate at a much slower rate than the population order *R.* The variable *ρ* is a key parameter that controls the degree of bistability in *R* time series. Under the influence of a weak *ρ* (*eq.* 4), the Kuramoto model could demonstrate dynamics resembling that of a *supercritical* stochastic Hopf bifurcation^25,34^ which is also controlled by a weak *ρ* (*eq.* 1). Specifically, by gradually increasing *κ*, a subcritical Kuramoto ensemble would reach a critical point, where the subcritical fixed point (quiescent state) loses stability and a smooth transition to a critical phase takes place. The time course of *R* in this classic critical scenario follows a Gaussian distribution^24–26^.

On the other hand, when the Kuramoto model is controlled by a high *ρ*, the ensemble would express bistable criticality in *R* that resembles the dynamics of a *subcritical* stochastic Hopf bifurcation. In this scenario, a seemingly quiescent ensemble suddenly shows supercritical hypersynchrony – before the quiescent fixed-point loses stability following further increases in *κ*. Thereby, the time course of *R* is characterized as bimodal, supporting erratic switching between low and high activity modes. Our modeling results confirmed this prediction. When *ρ* is held at high value, the pdf of *R* as a function of *κ* (the colour plot in top panel, Fig. 1E) coincides with the prediction of the first order phase transition with a moderate width of bistable regime (top, Supplementary Fig. 1A) – as comparing to theoretical possibility of a much wider bistable regime ^25,35^. When *ρ* is held at low value (bottom, Fig 1E), the model dynamics accords with the prediction of supercritical Hopf bifurcation as the classic criticality ^7,23^. When the coupling is too strong, the ensemble only dwells on the supercritical state of hypersynchrony. To better demonstrate the effect of *ρ* on R, we also simulated slowly fluctuating *ρ* with *κ* held constant and found that various waveforms of *ρ* can result in rich bistable dynamics in R, even when the temporal average of *ρ* was approximately the same (Supplementary Fig. 1B).

The model shows high degree of bistability under the influence of strong state-dependent noise, which results in a tendency to stay in either an up-state or a down-state thus avoiding moderate level of synchrony as expected in the classic criticality models. This up state corresponds to resource-demanding large amplitude oscillations (limit cycle), which eventually leads to depletion and ensuing returning of a low amplitude, subcritical down-state fixed point attractor for recovery. On the extreme spectrum of such bifurcation and the underlying slow variable is epileptic seizure ^52,53,104^.

#### SEEG cortical sampling statistics

Preprocessing of the SEEG data yielded 7,019 SEEG contacts in various cortical and subcortical gray matter locations (Supplementary Fig. 2A). For investigating the LRTCs and bistability of cortical dynamics that were tentatively considered as normal, contacts recorded from subcortical structures and epileptogenic zones were excluded. Contacts with more than 2.5% samples identified as “spiky” were also excluded (Supplementary Fig. 2A-B, see suppl. Methods). Thereby, 3/7 of available contacts were excluded, and in the resulted 4,142 contacts, a small fraction cannot be reliably assign a parcel by the segmentation software ^77^ and therefore were also excluded. This resulted in 4,122 contacts (66.8 ± 24.5 per patient, range: 4 to 123) for analyses. Although the cortical sampling was heterogeneous across patients, with 4,124 cortical nEZ contacts, we were able to cover 90 out of the 100 Schaefer parcels with each parcel sampled by at least 3 subjects and 10 contacts (see also Supplementary Fig 2C-D).

#### Narrow-band DFA and BiS estimates in MEG and SEEG

SEEG and MEG demonstrated differentiated spectral peaks and magnitude of DFA and BiS estimates (Fig 2). We speculated that these discrepancies between SEEG and MEG might be attributable to two factors (or the combination of both):

First, it was due to different brain states in healthy vs epileptic brains. High degree of bistability has been suggested as an early sign of shift to catastrophic events in ecosystems ^29^, considering the likely universal nature of bistability ^31,35^, high bistability outside of the visual system *(e.g.,* default model and limbic systems) in SEEG data could be a sign of transition to catastrophic events (*i.e.*, seizures) in the epileptic brain. It could be a great clinical interest to further this line of work to find solutions to reverse high bistability to mildly bistability or smooth phase transitions (*e.g.*, dampening) for eluding the catastrophic events in the brain ^105^. Moreover, epilepsy is known to affect brain rhythms especially the δ and θ band synchrony ^106,107^, which might be an explanation to the presence of high bistability in slow rhythms below α band in SEEG.

Second, mesoscopic SEEG recording is able to pick up highly local signals and thus more sensitive to specificity such as difference between functional systems, which MEG is unable to SEEG signals is highly localized to the sampled tissue surrounding the electrode contacts (2mm in length) and minimally affected by signal mixing ^108^, whereas MEG sensors are at least several centimeters away from the cortex, and its signals are therefore the linear summation of unknown number of cortical sources. According to the central limit theorem, even though individual processes are non-Gaussian, the linear combination of multiple such processes will appear Gaussian.

#### Topological similarity between LRTCs and bistability

As a preliminary inspection of topological similarity, all-to-all correlations were computed indiscriminately between group-level parcel DFA and BiS estimates, between MEG and SEEG, and across all frequencies. Within metrics, well delineated clusters of a slow and a fast frequency band were observed (along diagonal, Supplementary Fig. 6). There were positive correlations between DFA and BiS in both MEG and SEEG data, and some negative correlations between MEG and SEEG. The frequency-collapsed θ–α and γ-band DFA and BiS estimates were next inspected (Fig. 3C). The similarity between band-clustered cortical maps of 100 Schaefer parcels (Supplementary Fig. 7) converged with narrow-band observation (Supplementary Fig. 6). On systems-level (Supplementary Fig. 8), subjects’ band-clustered DFA and BiS were correlated in almost all functional systems except for SEEG θ–α band limbic system (Spearman’s rank correlation, *p* < 0.01).

#### Anatomical specificity of bistability and LRTCs

While there were no systems-wise differences in MEG, SEEG θ–α band BiS and DFA estimates, γ band DFA estimates between Yeo functional systems were different (Kruskal-Wallis test, *p* < 0.05, Fig. 3F).

In MEG, the mean θ–α and γ-band DFA and BiS estimates of Yeo systems appeared similar (thick black lines, Supplementary Fig. 9A-B), confirming high correlations between DFA and BiS estimates observed in both group and systems-level in individuals (Supplementary Fig. 7-8, respectively). With these source-modeled MEG data, we replicated θ–α band resting-state bistability in visual areas that was reported previously in EEG sensors ^26,109^. Furthermore, the visual (VIS) and somatosensory (SM) systems showed higher BiS and DFA, whereas fronto-parietal (FP) and default-model (DEF) systems showed lower BiS and DFA estimates than the null-hypothesis observations (line plots, Supplementary Fig 9A). The surrogate data of no systems-wise differences were constructed by shuffling Yeo system labels of the parcels. In MEG γ-band, DFA estimates were similar in magnitude to that of θ–α band, but BiS estimates were about half of the magnitude of θ–α band. In particular, the dorsal-attention (DAN) and the limbic (LIM) systems showed the highest γ-band DFA, whereas VIS had the lowest DFA. There were no differences in DFA and BiS estimates between Yeo systems (Wilcoxon’s signed-rank test, alpha@0.05, FDR corrected) confirming the negative findings of the Kruskal-Wallis tests (Fig. 3F).

In SEEG, the θ–α and γ band mean DFA and BiS estimates of Yeo systems appeared different (Supplementary Fig. 9 C-D), which confirmed low correlations in group and systems-level in individuals (Supplementary Fig. 7 & 8). Although visual systems in SEEG and MEG data were comparable in θ–α band BiS and DFA estimates, in SEEG, the visual system showed lowest θ–α band BiS and DFA estimates.

The post-hoc tests (unpaired t-test, *p* < 0.05, FRD corrected) for between-system differences in DFA and BiS estimates (see also Fig. 3F) revealed that DAN showed higher θ–α band DFA, FP and DEF showed higher θ–α band BiS, ventral-attention (VAN) showed lower γ band BiS than most of other systems (interaction matrices, Supplementary Fig. 9 C-D). On the other hand, the VIS and VAN show low DFA estimates. System-wise difference in DFA was only observed between LIM and VAN, which validates the negative finding of systems’ effect on mean DFA (γ band pink dot in Fig 3F). On the other hand, the VIS, DAN, and LIM system showed high γ band BiS estimates, and VAN and FP systems have low BiS estimates. The VIS system had highest γ band BiS estimates.

#### Classifying epileptogenic zones (EZ)

Across subjects, there was large variability in the number of EZ contacts, *i.e.,* the target variables of the classifier (Supplementary Fig. 11A). To ensure there were enough data for the classifier within subjects, a selection criterion was imposed such that each patient should have at least five EZ and five nEZ contacts (red dashed lines, Supplementary Fig. 11A) and at least a total of 30 contacts. Thus, 55 subjects met these criteria and were selected for the classification analysis. On average, each subject had 28.5±17.0 (mean, std) EZ and 66.4 ±21.2 nEZ, and on population the ratio of EZ:nEZ = 1/2.3 with some variability across Yeo systems, among which subcritical contacts were ten times more likely to be EZ than nEZ (Supplementary Fig. 11B).

On population level, between EZ and nEZ contacts, several bands showed differences in normalized DFA and BiS estimates (Fig. 5A-C). However, we were more interested in classifying EZ contacts in individuals. As a liability check, we classified pooled individual contacts using all nine features, *i.e.*, δ, θ–α, β, and γ band DFA and BiS estimates plus SEEG contact loci in Yeo systems. The classification was performed using the random forest algorithm^39^ with a total of 5,217 contacts from 55 patients (Supplementary Fig. 11B) with randomly split 20% and 80% as testing and training set, respectively. The analysis of the receiver operating characteristics (ROC) of classification outcome revealed an area under the curve (AUC) of 0.8, and thus confirming useful information among these features for within individual classification.

Global and local feature importance to the random forest classifier were next assessed with SHAP values. The results supported the hypotheses that γ- and β-band BiS, contact-locus and δ-band DFA were indeed the most important features to tell EZ apart from nEZ contacts (Fig. 5D). To better understand the impact of the features on the classification outcome, the within-subject EZ classifications were carried out with four incremental feature sets as using *i*) DFA alone, *ii*) BiS alone, *iii*) combing DFA and BiS, *iv*) combing DFA, BiS, and contact-locus in Yeo systems.

The overall classification outcome was variable across Yeo systems and classifying EZ contacts in limbic systems yielded best outcome (large AUC, Supplementary Fig. 11 D). Using DFA alone, the algorithm performed poorly (gray line, Supplementary Fig. 11E). The individual subject ROC curves showed large variability (Supplementary Fig. 11 F-I), and overall combing DFA, BiS, and contact-locus yielded best outcome (black curve, Supplementary Fig. 11J). Last, the mean AUC of ROC in Yeo systems (Supplementary Fig. 11E) and in individual patients (Supplementary Fig. 11J) were compared against 1,000 label-shuffled surrogate data (Fig. 5E).

### Supplementary (S.) Figs

**Supplementary Fig. 1.**
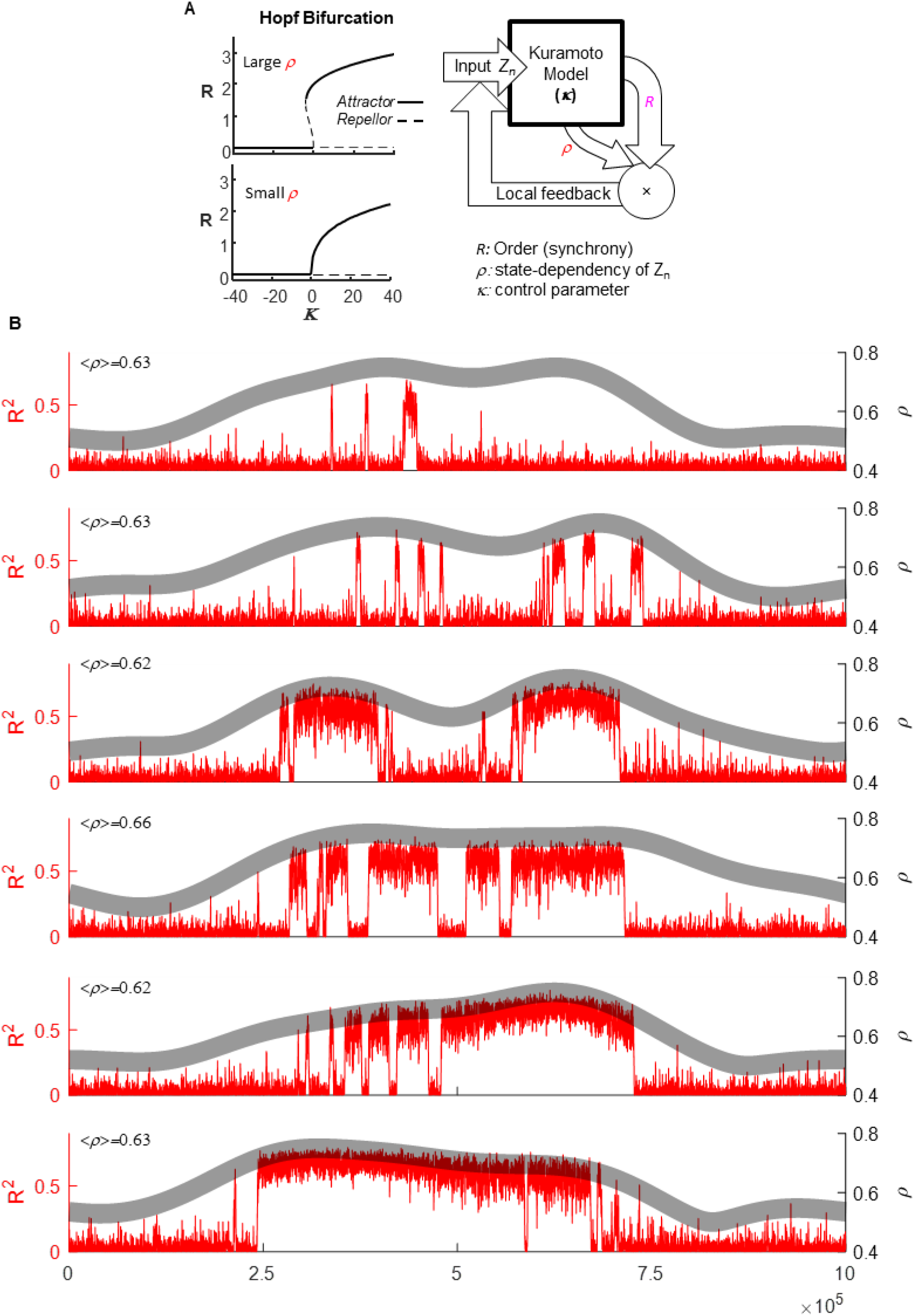
Fluctuations at two different time-scale in Kuramoto Model. (**A**) With coupling strength (*κ*) held constant and just below the critical point (see Fig 1), slow fluctuations in *ρ* (the thick gray band) result in (**B**) diverse patterns of bistability in the fast fluctuating mean field of the Kuramoto model (red).

**Supplementary Fig. 2.**
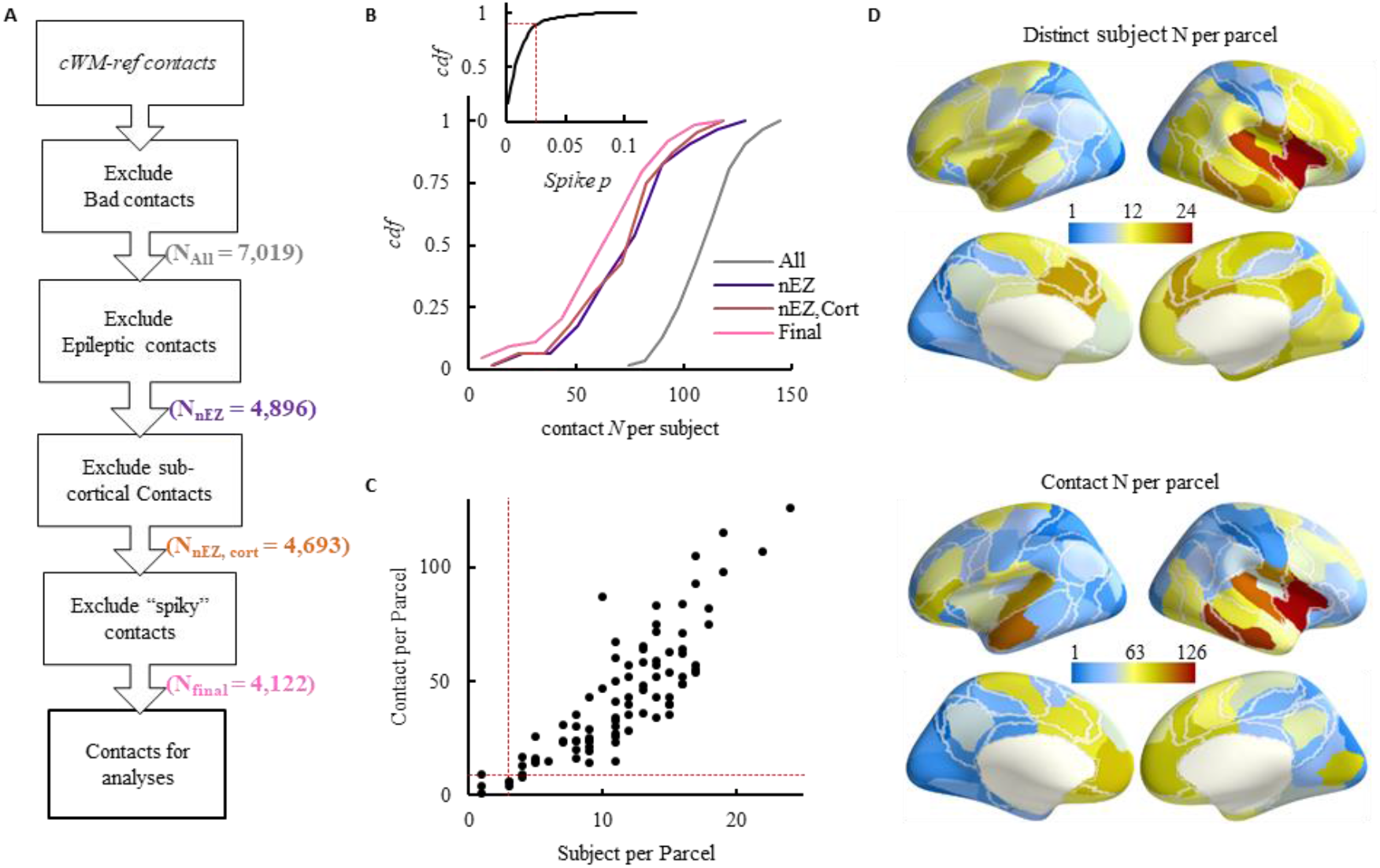
SEEG electrode selection criteria and population sampling statistics. (**A**) SEEG contact selection criteria, in brackets are the number of contacts after a specific criterion was applied. (**B**) The cumulative distribution function (*cdf*) of contact number per subject after specific criteria were applied; colour code is the same as in (**A**); inset is the *cdf* of spike samples out of 10 min resting from 4,693 SEEG contacts, red dashed line indicates the threshold in box 5 in (**A**). (**C**) Contact number as a function of number of distinct subjects per parcel (after morphing SEEG contacts into Schaefer 100-parcels); one marker represents one Schaefer 100-parcle, red dashed lines are exclusion criteria, *i.e.,* at least 3 subjects and 10 contacts per parcel. (**D**) Visualization of distinct subject number and SEEG contact number per Schaefer parcel as shown in (**C**). cWM-ref: closest white-matter contact reference scheme (Arnulfo et al., 2015).

**Supplementary Fig. 3.**
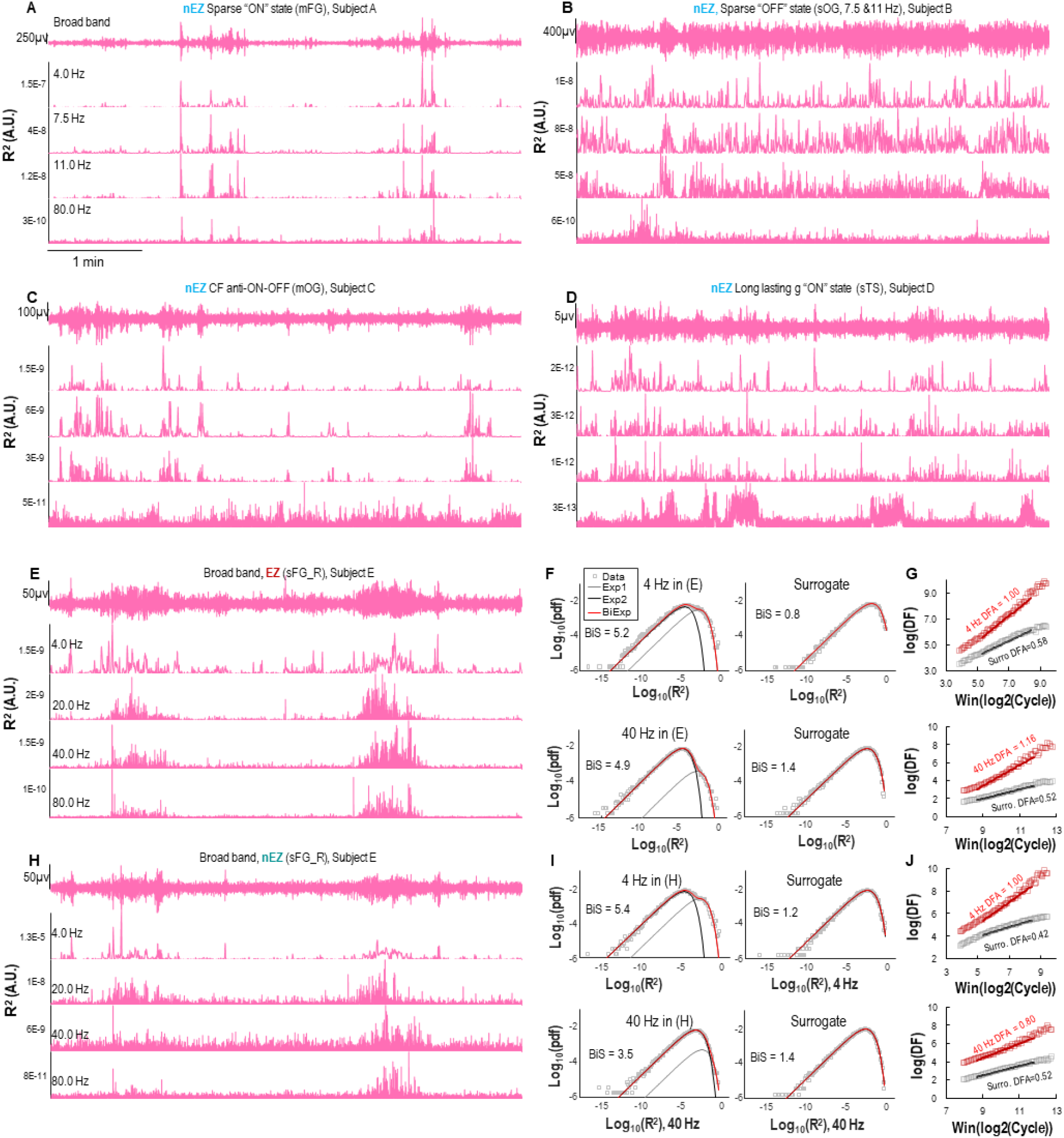
SEEG time series, bistability and DFA fitting. (**A–D**) Examples of bistability in nEZ SEEG contact signals from four distinct subjects. mFG: middle frontal gyrus; sOG: superior occipital gyrus; mOG: middle occipital gyrus; sTS: superior temporal sulcus. (**E-J**) Examples of bistable time series and model fitting of an EZ (**E**) and a nEZ (**H**) contact. These two contact locations were 19.7 mm apart and recorded with two distinct electrodes from the superior frontal gyrus (sFG) and were referenced with the same nearest white matter contact (Arnulfo et al., 2015). (**F**) Examples of bi-exponential model fitting for BiS estimates and (**G**) DFA power-law fitting of 4 Hz and 40 Hz narrow-band real and surrogate time series of the EZ contact from (**E**). (**I**) Bi-exponential fitting and (**J**) DFA power-law fitting of 4 Hz and 40 Hz narrow-band real and surrogate time series of the nEZ contact from (**H**). The DFA fitting plot reads as, when the observation window size doubles (by narrow-band cycle length – irrespective of frequency), the detrended fluctuation increase by a constant rate of log(DF).

**Supplementary Fig. 4.**
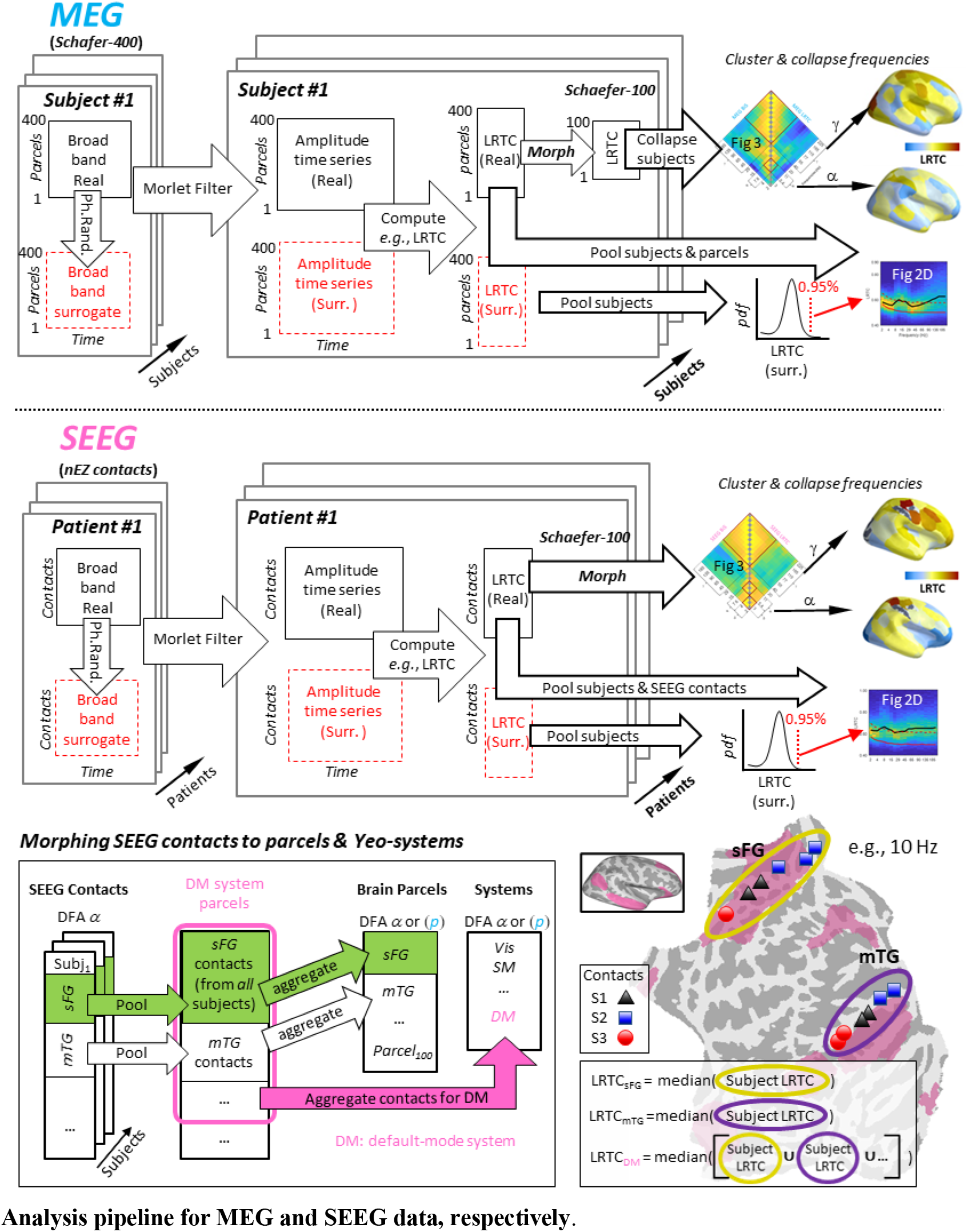
Analysis pipeline for MEG and SEEG data, respectively.

**Supplementary Fig. 5.**
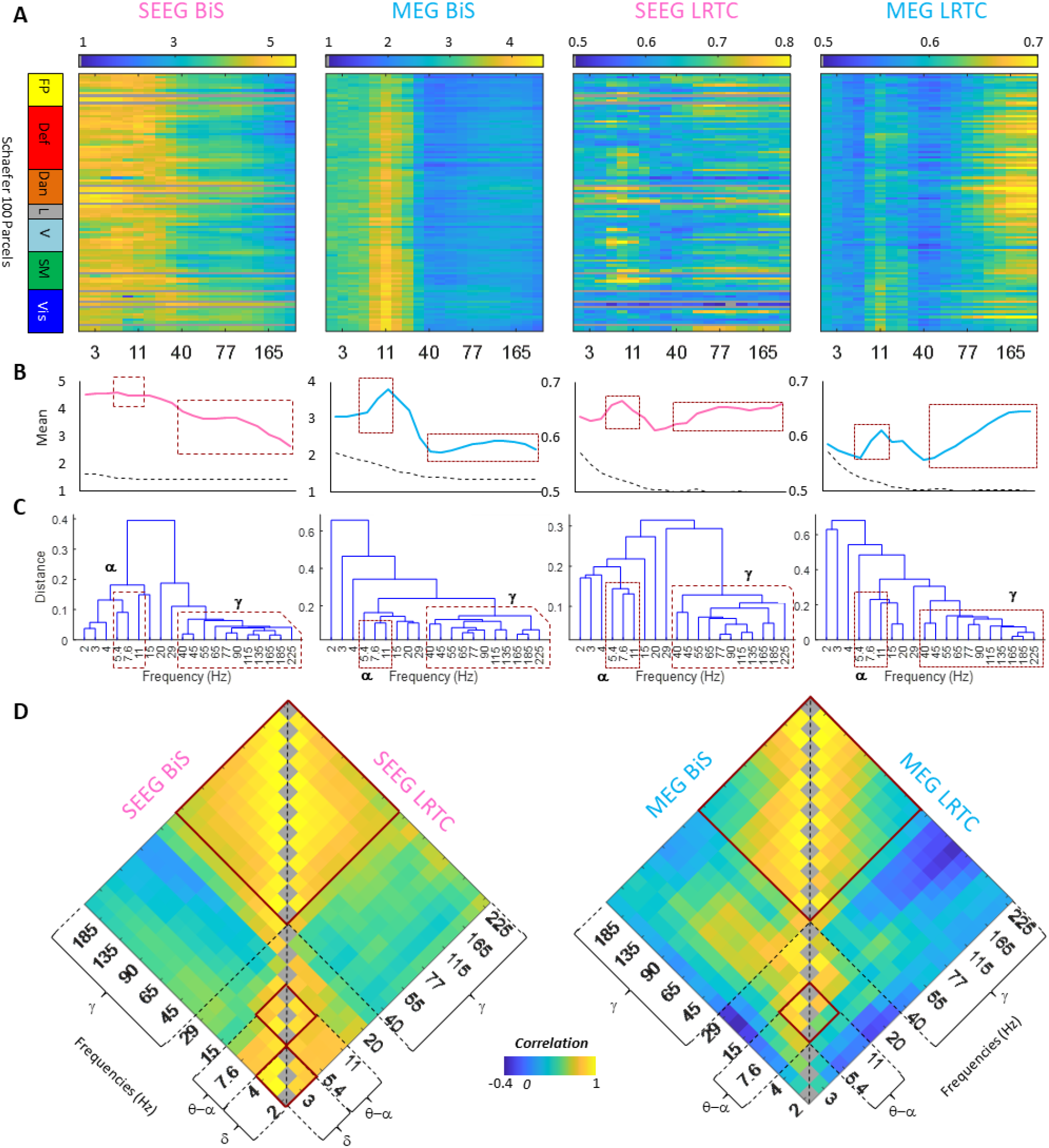
The cortical origins of DFA and BiS are similar between neighbouring frequencies but different between slow and fast rhythms demarcated at 40 Hz. (**A**) BiS and DFA of each Schaefer parcel (y axis) observed at narrow-band frequency (x axis) ranging from 2 to 225 Hz for SEEG and MEG data. The 10 gray-out rows are the excluded SEEG parcels due to under sampling. (**B**) Parcel mean across frequencies of real data (solid lines) and surrogate (dashed lines). (**C**) The distance between each frequency’s spatial similarity. (**A**-**C**) share the same *x*-axis (Frequency); (**D**) Cross-frequency adjacency matrices of topological similarity, the same as Fig 3A; correlation is the Spearman’s *r* between Schaefer Atlas parcels’ DFA or BiS of two different frequencies.

**Supplementary Fig. 6.**
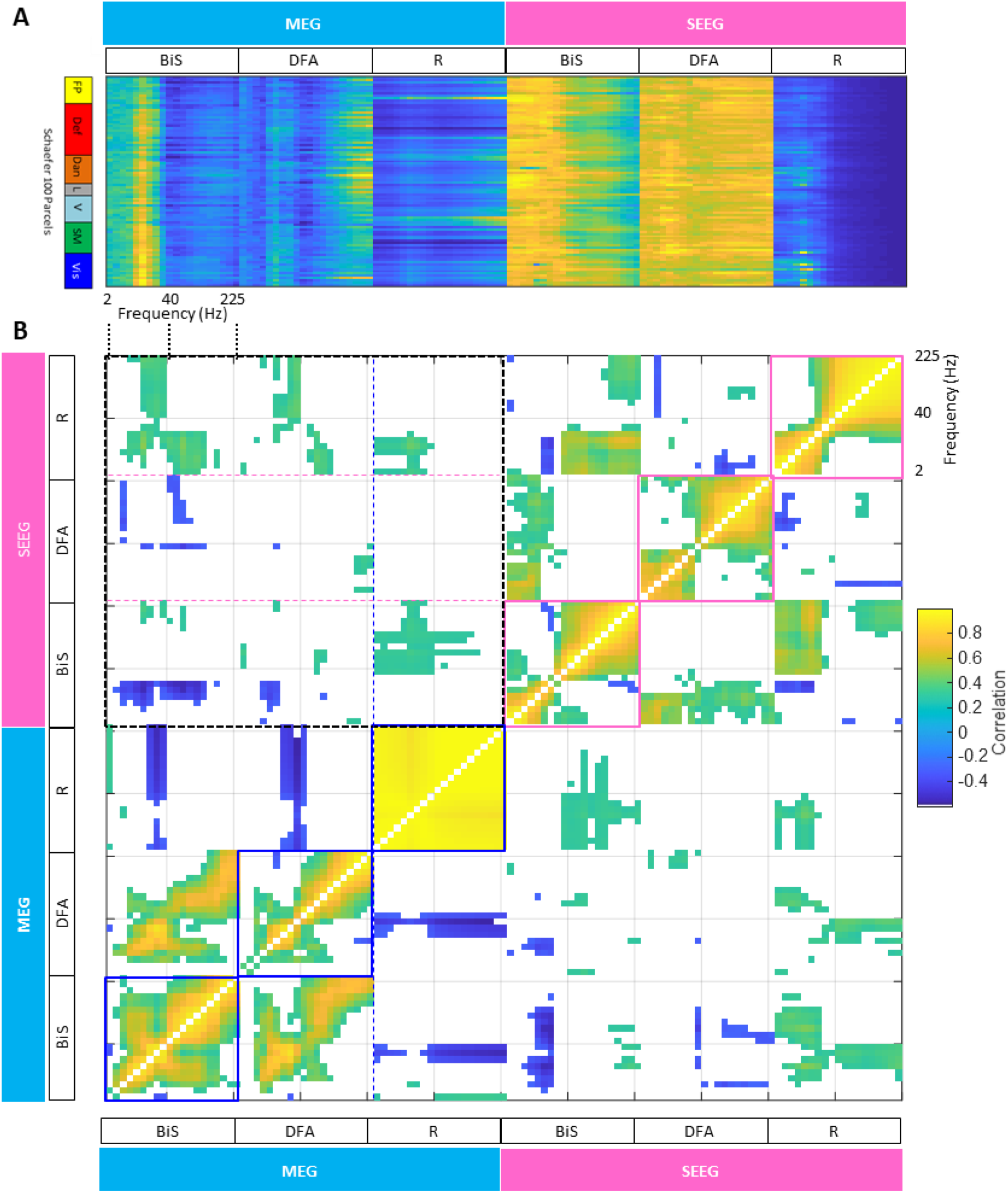
All-to-all cross correlations between frequencies, metrics and datasets. (**A**) Normalized group-level Schaefer 100-parcel metrics afo frequencies. Normalization y = (x(i)-min(x)) /(max(x)-min(x)). (**B**) All-to-all topological correlations between R, DFA, and BiS and between MEG and SEEG data (Spearman’s rank order *r, p*>0.01, not controlled for FDR).

**Supplementary Fig. 7.**
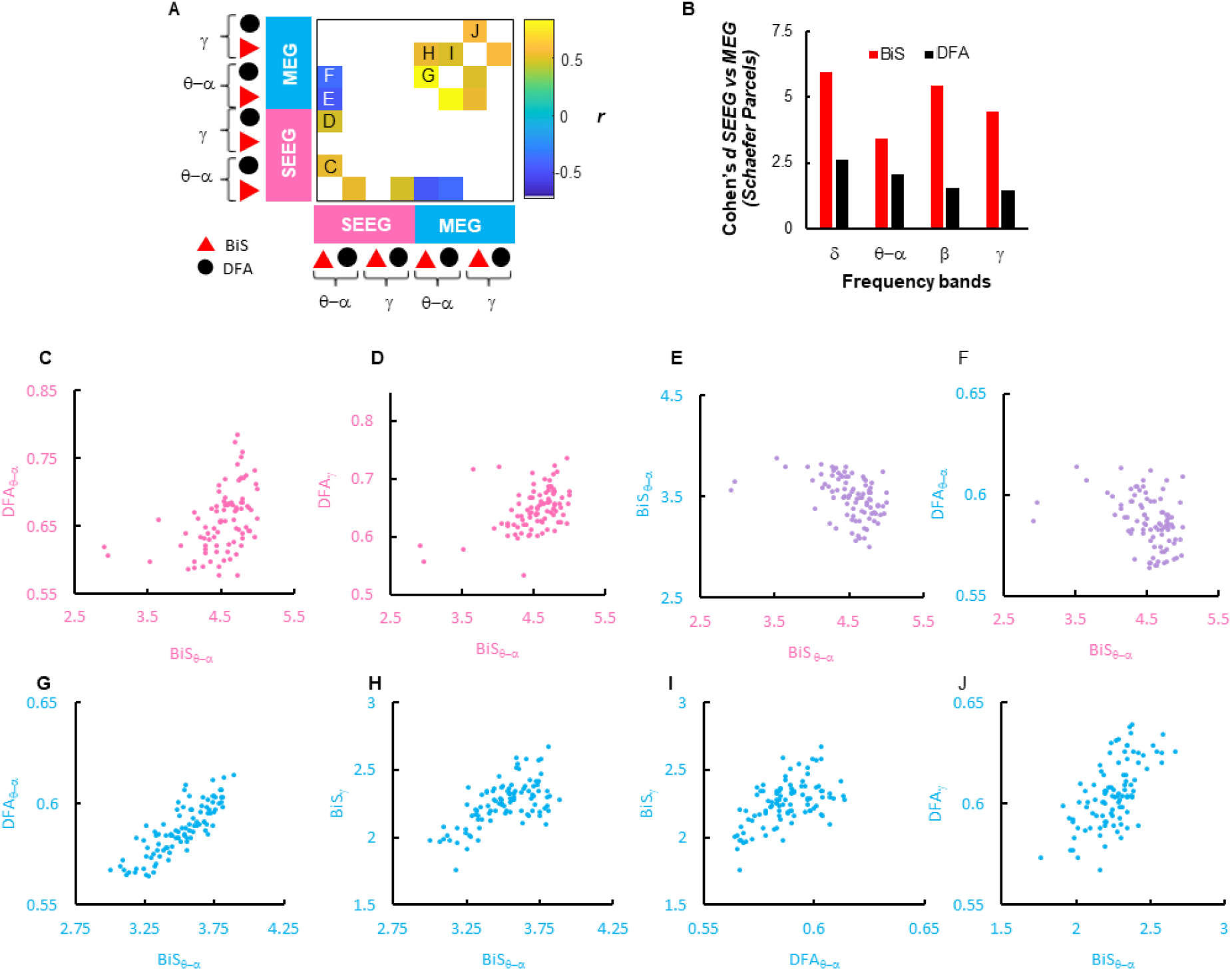
DFA and BiS estimates correlated in MEG and SEEG cortical parcels. (**A**)The adjacency matrix of significant Spearman’s correlation between band-collapsed DFA and BiS (p < 10^-6^, FDR corrected). (**B**) The effect size of the differences between DFA and BiS of MEG and SEEG data in Schaefer parcels (n=90, due to exclusion of 10 SEEG parcels). (**C**-**J**) Scatter plots showing correlation between DFA and BiS estimates in band-clustered all-to-all correlation matrix (top); each data point corresponds to the group average metrics in one Schaefer 100-parcel (NMEG=100; NSEEG = 90).

**Supplementary Fig. 8.**
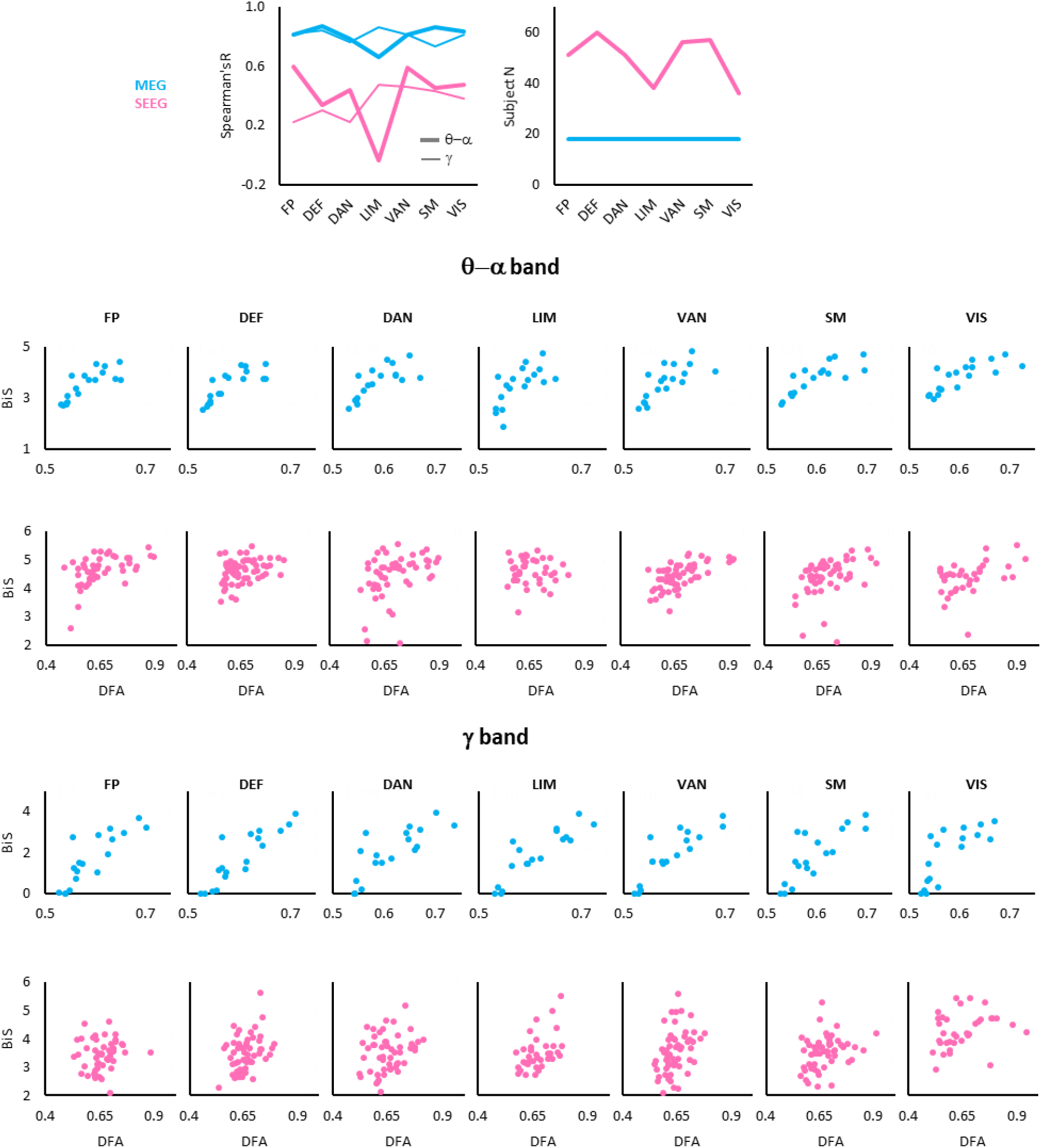
DFA and BiS are correlated in MEG and SEEG subsystems. Top left panel is from Fig 3H; top right is the number of subjects observed in each Yeo system: MEG N=18, and SEEG N=50±9, range: 36-60, variable subject N per system in SEEG due to heterogeneous spatial sampling. In scatter plots, each dot corresponds to the observation from one subject.

**Supplementary Fig. 9.**
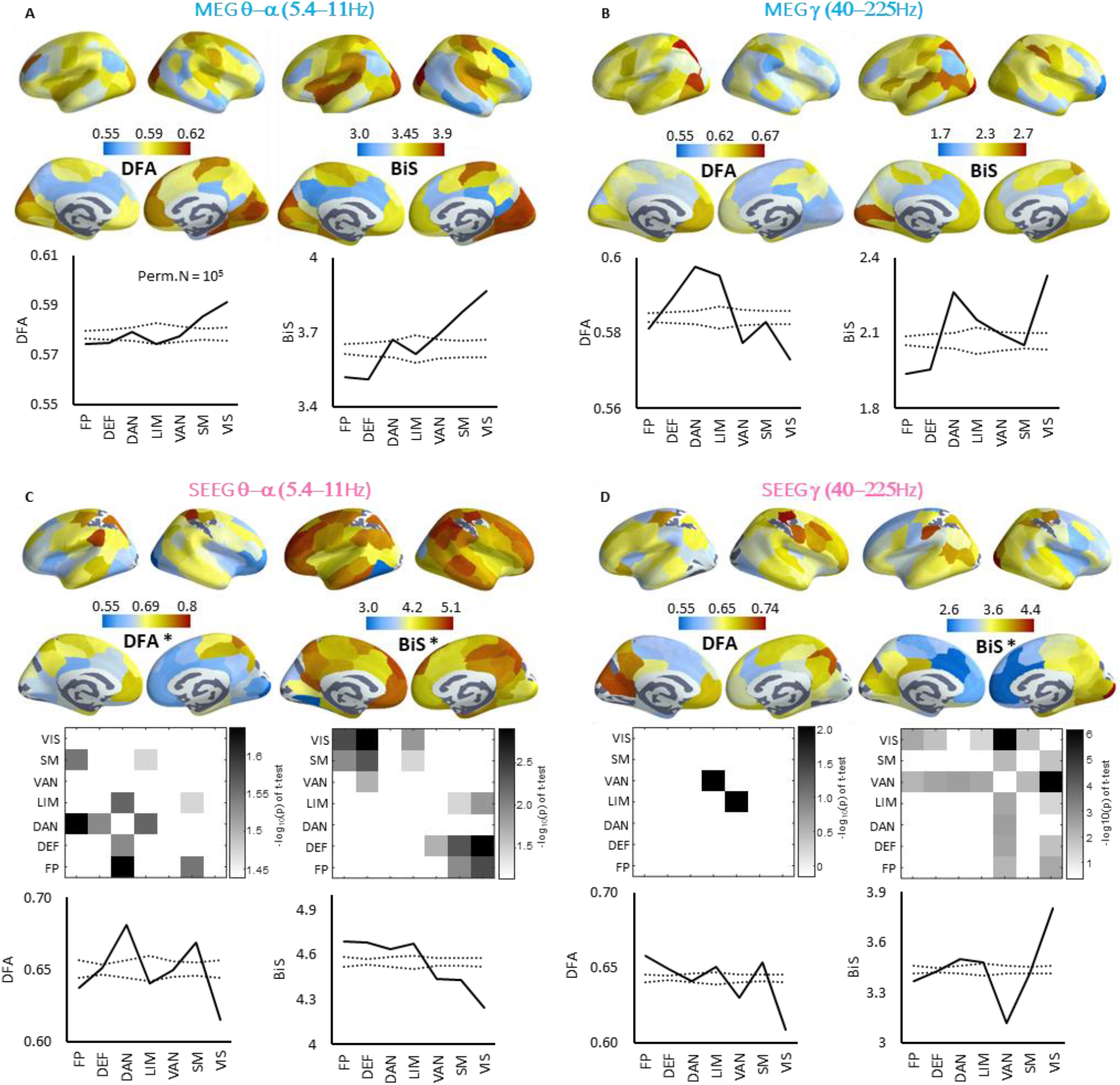
Systems-level differences and anatomical localization of DFA and BiS. These are post hoc tests for differences in DFA and BiS between Yeo systems (Fig 3I). (**A-D**) Neuro-anatomical localization of frequency-collapsed BiS and DFA; marker (*) in colour-bar label indicates significant differences between systems (Fig 3I). The line plots below the brains are median DFA/BiS of Yeo systems, dashed lines are 5%- and 95%-tile of permutation median (N_permutation_ = 10^5^). The adjacency matrices are the significant – log_10_(*p*) values of pairwise tests for between-system differences (SEEG: unpaired t-test; MEG: Wilcoxon’s signed-rank test, *p*<0.05, FDR corrected).

**Supplementary Fig. 10.**
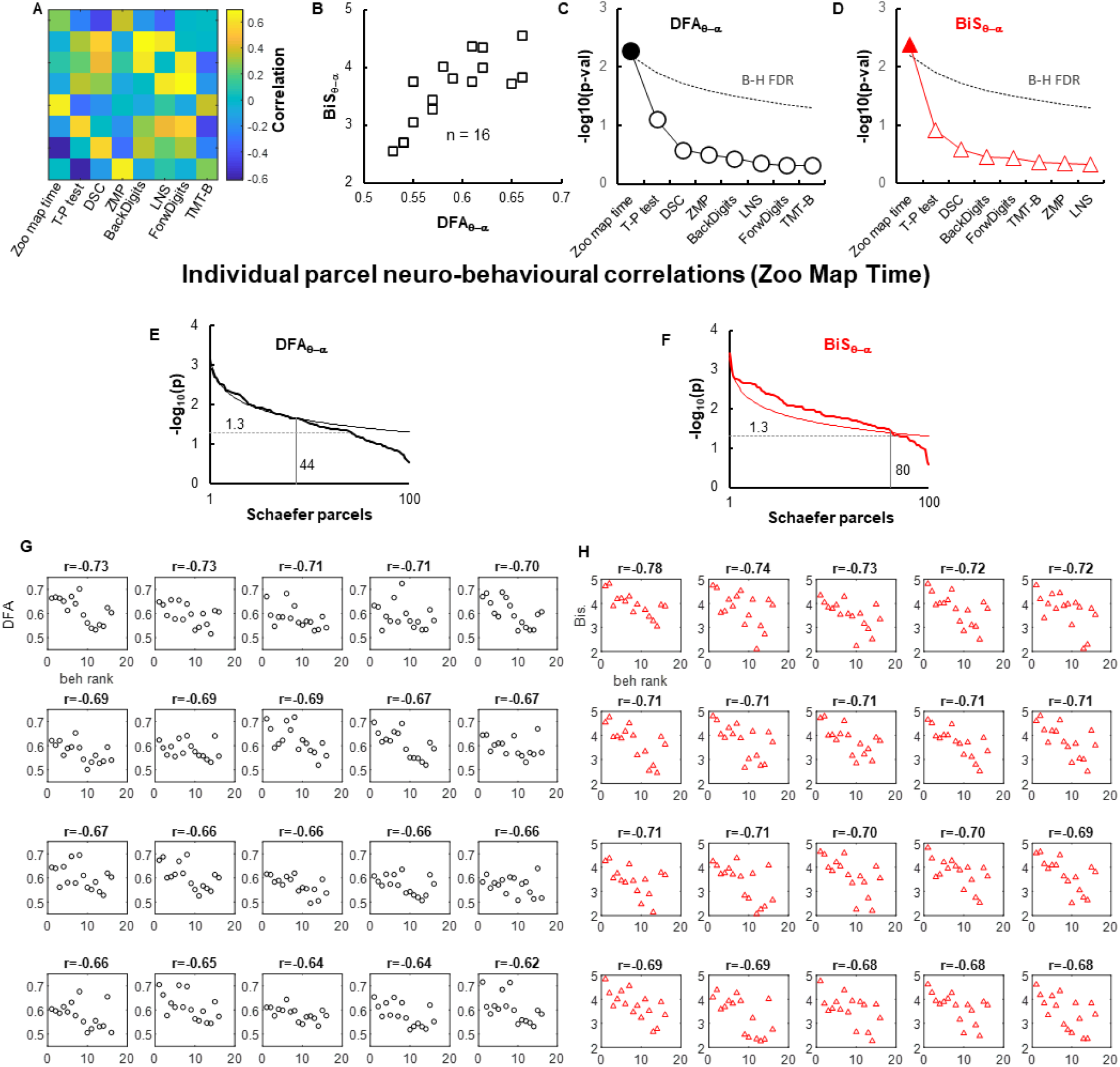
Behaviourally relevant DFA and BiS in MEG. (**A**) Left: percentage of parcels showed significant correlation between executive function score (normalized zoo map time) and parcel DFA and BiS; right: θ–α band BiS as a function of DFA. (**B**) The behavioural correlations (primary y-axis) and probability of parcels showing significant correlations (secondary y-axis) on functional systems-level (corrected for FDR). (**C**) Top row: cortical maps of behavioural correlations; bottom row: subject neuronal estimates as a function of executive score from exemplary regions of interest where each marker is the observation from one subject (**C-D**) The corresponding *p* values of correlation tests in Fig 4A and C, respectively; dashed lines indicate FDR (Benjamini-Hochberg procedure with FDR at 0.05 for 8 neuropsychological scores) adjusted *p* values for eight neuropsychological tests. (**E**) Sorted p-values (thick line) of individual parcel behavioural correlation for θ–α band DFA and (**F**) BiS estimates. Thin lines indicate Benjamini-Hochberg procedure with FDR at 0.05 adjusted *p* values. (**G**) Examples of parcel-level behavioural correlations to Zoom map time rank for a-band DFAs and (**H**) θ–α band BiS estimates, where each marker indicates one subject.

**Supplementary Fig. 11.**
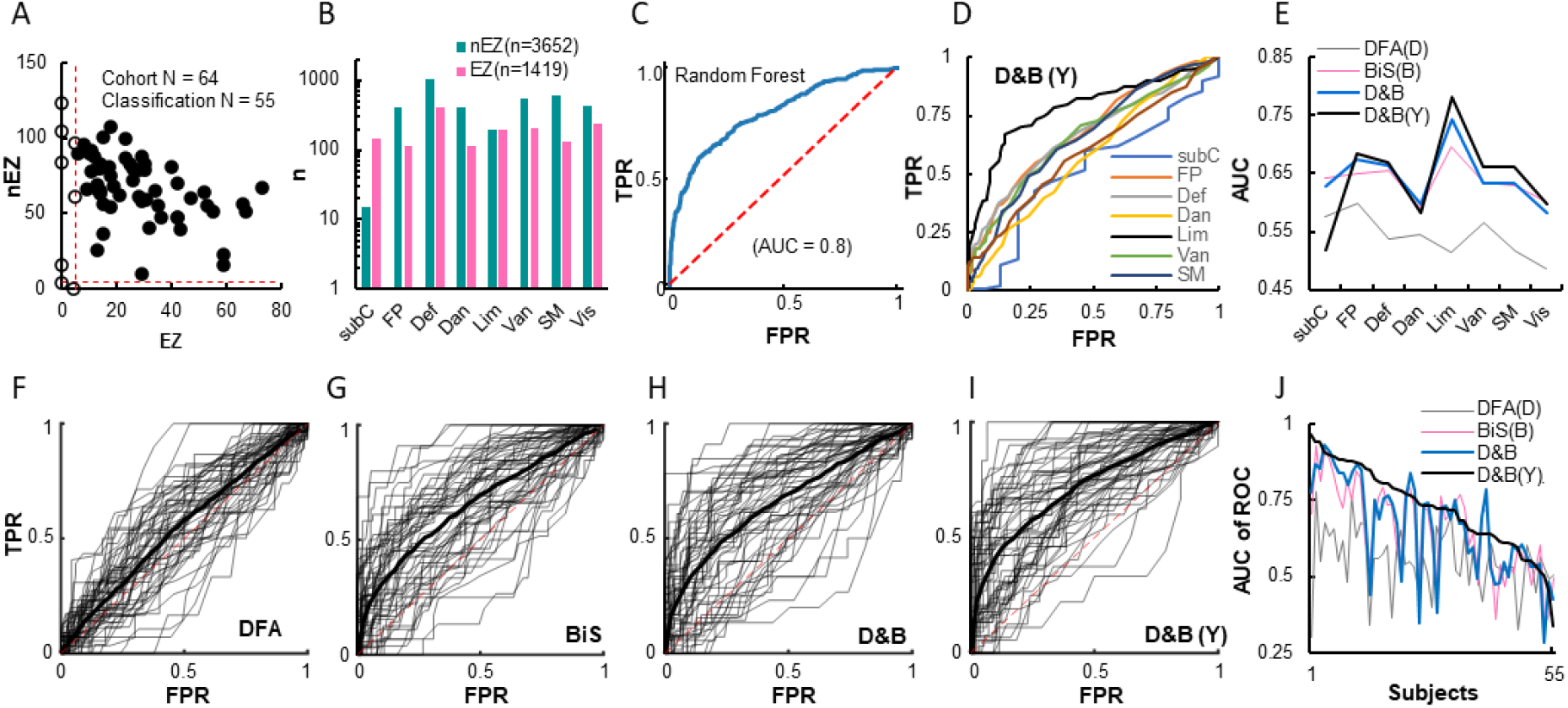
Classification of EZ and nEZ SEEG contacts (both cortical and subcortical) using band clustered DFA and BiS estimates. (**A**) Number of EZ and nEZ contacts in each subject; solid dots indicate subjects who met selection criteria for classification analysis; red dashed lines indicate selection criterion: each subject must have great than five EZ and nEZ contacts. (**B**) EZ and nEZ contacts located in Yeo’s systems and subcortical (subC) regions, data pooled over 55 subjects from (**A**). (**C**) An example of the receiver operator curve (ROC) of preliminary population-level classification using all contacts and all features with randomly split test and train set (20% and 80%, respectively); TPR: true positive rate; FPR: false positive rate; AUC: area under the ROC curve. (**D-J**) Leave-one(subject)-out classification results using random forest algorithm. (**D**) The ROC of each subsystem, classification with the full feature set, D&B(Y). (**E**) The AUC yielded from classification using DFA only, BiS estimates only, combining DFA and BiS (D&B), and full feature set D&B(Y), *i.e.,* DFA, BiS and SEEG contact loci. (**F-I**): individual (thin lines) and group average ROC (thick) yielded from classification using varying feature sets. (**J**) The AUC of ROCs shown in (**F-J**), subjects were sorted by the area under the ROC curve of D&B (Y) feature set.

## Notes

### Competing Interest Statement

The authors have declared no competing interest.

